# Distinct effects of CDK8 module subunits on cellular growth and proliferation in *Drosophila*

**DOI:** 10.1101/2024.04.30.591924

**Authors:** Xiao Li, Mengmeng Liu, Yue Xing, Ye Niu, Tzu-Hao Liu, Jasmine L. Sun, Yanwu Liu, Rajitha-Udakara-Sampath Hemba-Waduge, Jun-Yuan Ji

## Abstract

The Mediator complex, composed of about 30 conserved subunits, plays a pivotal role in facilitating RNA polymerase II-dependent transcription in eukaryotes. Within this complex, the CDK8 kinase module (CKM), comprising Med12, Med13, CDK8, and CycC (Cyclin C), serves as a dissociable subcomplex that modulates the activity of the small Mediator complex. Genetic studies in *Drosophila* have revealed distinct phenotypes of CDK8-CycC and Med12-Med13 mutations, yet the underlying mechanism has remained unknown. Here, using *Drosophila* as a model organism, we show that depleting CDK8-CycC enhances E2F1 target gene expression and promotes cell-cycle progression. Conversely, depletion of Med12-Med13 affects the expression of ribosomal protein genes and fibrillarin, indicating a more severe reduction in ribosome biogenesis and cellular growth compared to the loss of CDK8-CycC. Moreover, we found that the stability of CDK8 and CycC relies on Med12 and Med13, with a mutually interdependent relationship between Med12 and Med13. Furthermore, CycC stability depends on the other three CKM subunits. These findings reveal distinct roles for CKM subunits *in vivo*, with Med12-Med13 disruption exerting a more pronounced impact on ribosome biogenesis and cellular growth compared to the loss of CDK8-CycC.

**Significance:** The CDK8 kinase module (CKM), comprising CDK8, CycC, Med12, and Med13, is essential in the Mediator complex for RNA polymerase II-dependent transcription in eukaryotes. While expected to function jointly, CKM subunit mutations result in distinct phenotypes in *Drosophila*. This study investigates the mechanisms driving these differing effects. Our analysis reveals the role of Med12-Med13 pair in regulating ribosomal biogenesis and cellular growth, contrasting with the involvement of CDK8-CycC in E2F1-dependent cell-cycle progression. Additionally, an asymmetric interdependence in the stability of CDK8-CycC and Med12-Med13 was observed. CKM mutations or overexpression are associated with cancers and cardiovascular diseases. Our findings underscore the distinct impacts of CKM mutations on cellular growth and proliferation, advancing our understanding of their diverse consequences *in vivo*.

## Introduction

Transcriptional activators recruit various multiprotein complexes to orchestrate precise control over RNA polymerase (Pol II) initiation, elongation, and termination (1–3). Among these transcriptional complexes, the Mediator complex, the largest protein complex, plays a critical role and comprises approximately 30 conserved subunits in eukaryotes, categorized into four modules: the head module, the middle module, the tail module, and the CDK8 kinase module (CKM) (2, 4–7). The CKM complex consists of four subunits, CDK8 (or its paralog CDK19, also known as CDK8L, in vertebrates), CycC, Med12 (or MED12L in vertebrates), and Med13 (or MED13L in vertebrates). Two prevailing models have been proposed to explain how CKM regulates transcription. In one model, CKM can reversibly associate with the small Mediator complex, which comprises the head, middle, and tail modules. This association blocks the interactions between the small Mediator complex and Pol II, leading to the inhibition of the Pol II-dependent transcription (8–10). An alternative but nonexclusive model suggests that the CDK8 kinase within CKM can phosphorylate a variety of transcription factors, thus modulating their transcriptional activities in various biological contexts (11–21).

Biochemical purification of the CKM in yeast and mammals have showed that the four CKM subunits can be purified together, and the structure of CKM has been determined through the cryogenic electron microscopy (cryo-EM) (9, 22–24). In addition, both CycC and Med12 are required for the *in vitro* kinase activity of CDK8 (22). Notably, direct interactions between the N-terminal segment of MED12 and the T-loop of CDK8 are pivotal for CDK8 activation (23, 24). Consistent to these findings, missense mutations located in exon 2, which encodes a portion of the N-terminal segment of MED12, have been identified in over 62% of uterine leiomyomas worldwide (25). These mutations significantly reduce the kinase activities of CDK8-CycC or CDK19-CycC (26). Collectively, these biochemical analyses suggest that the four subunits of the CKM complex operate together as a unified entity to fulfill their biological functions.

Despite these comprehensive biochemical and structural studies, genetic analyses in *Drosophila* have revealed that the four subunits of CKM complex may have distinct roles during development. For instance, null mutations in *kto* (encoding Kohtalo, the *Drosophila* Med12 ortholog) and *skd* (encoding Skuld, the *Drosophila* Med13 ortholog) result in embryonic lethality, while null mutations in *cdk8* and *cycC* lead to pupal lethality (27–29). In addition, genetic ablation of Med12 (Kto) or Med13 (Skd) in *Drosophila* eye or leg imaginal discs leads to severe defects in adult eyes and legs, whereas loss of CDK8 or CycC in eye or leg discs causes only minor leg defects and no eye defects (27, 30). These distinct phenotypic outcomes challenge the assumption, suggested by the biochemical and structural studies, that the four CKM subunits function equally as an indivisible CKM complex. Mutation or amplification of the CKM subunits have been identified in various cancers and cardiovascular diseases (31–33), underscoring the importance of understanding the cellular consequences of these mutations *in vivo* and the underlying mechanisms.

In this study, we investigated the mechanism underlying the distinct *in vivo* functions of CDK8-CycC and Med12-Med13 in *Drosophila*. We generated a series of transgenic *Drosophila* lines enabling individual or combinatorial depletion of the four CKM subunits. Our genetic analyses confirmed distinctive phenotypes resulting from the depletion of these CKM subunits in *Drosophila* eyes and wings. At the cellular level, we observed that depleting CDK8-CycC promoted cell proliferation, while depleting Med12-Med13 did not noticeably affect cell proliferation. These observations are consistent with the upregulation of E2F1 target genes when CDK8-CycC was knocked down, whereas depleting Med12-Med13 primarily affected the expression of genes involved in ribosomal biogenesis and cellular growth. Furthermore, we observed that the stability of the CDK8-CycC pair and the MED-Med13 pair is asymmetrically interdependent, with a more substantial impact of the loss of MED12-Med13 on the entire CKM than the disruption of CDK8-CycC. Considering reports demonstrating mutations or amplifications in the four CKM subunits in various human diseases such as cancers and cardiovascular disorders (31–34), elucidating the specific consequences of these mutations on CKM function, as opposed to the kinase activities of CDK8 or CDK19, may advance our understanding of how mutations in distinct CKM subunits contribute to tumorigenesis and other pathological contexts.

## Results

### Depletion of the CKM subunits distinctively impacts *Drosophila* eye development

Previous studies demonstrate that null mutants of *Cdk8* and *CycC* result in pupal lethality, while null mutants of *Med12* (*kto*) and *Med13* (*skd*) lead to embryonic lethality (27–30). Investigating the specific roles of individual CKM subunits during development is crucial but technically challenging, particularly in simultaneously targeting these subunits in various combinations in a tissue-specific manner. To overcome this challenge, we utilized the pNP vector, which enables the concurrent expression of multiple shRNAs that specifically target different genes (35). We generated a series of transgenic RNAi *Drosophila* lines using the pNP vector to deplete the four CKM subunits individually or in all conceivable combinations (Table S1). Specifically, we created four lines for depleting each of the four subunits independently (*UAS-CDK8-RNAi*, *UAS-CycC-RNAi*, *UAS-Med12-RNAi*, and *UAS-Med13-RNAi*). Additionally, six lines were designed to target various combinations of the two subunits of the CKM (*UAS-CDK8-i-CycC-i*, *UAS-CDK8-i-Med12-i*, *UAS-CDK8-i-Med13-i*, *UAS-CycC-i-Med12-i*, *UAS-CycC-i-Med13-i,* and *UAS-Med12-i-Med13-i*). Furthermore, four lines were created for the simultaneous depletion of three subunits (*UAS-CDK8-i-CycC-i-Med12-i, UAS-CDK8-i-CycC-i-Med13-i, UAS-CDK8-i-Med12-i-Med13-i*, and *UAS-CycC-i-Med12-i-Med13-i*). Lastly, one line was established for the simultaneous depletion of all the four subunits (*UAS-CDK8-i-CycC-i-Med12-i-Med13-i*) (see Materials and Methods for details). To ensure consistent expression levels of shRNAs, all these constructs were site-specifically integrated into the *attP2* site on the third chromosome (36).

To analyze the *in vivo* function of distinct CKM subunits, we used *eyeless-Gal4* (*ey-Gal4*) to deplete these subunits in various combinations in *Drosophila* eye. Compared to the control (Fig. 1A), depleting CDK8 alone (Fig.1B), CycC alone (Fig. 1C), or both CDK8 and CycC simultaneously (Fig. 1D) had no discernible effects on *Drosophila* eye development. However, depleting Med12 (Fig. 1E) or Med13 (Fig. 1F) resulted in a significant reduction in the size of the adult eyes, as quantified in Fig. 1Q. These observations are consistent to a previous report showing that eye-specific removal of CDK8 or CycC did not affect eye development, whereas the removal of Med12 alone, or Med12 in combination with CDK8 led to a substantial reduction in eye size (27).

**Fig. 1.**
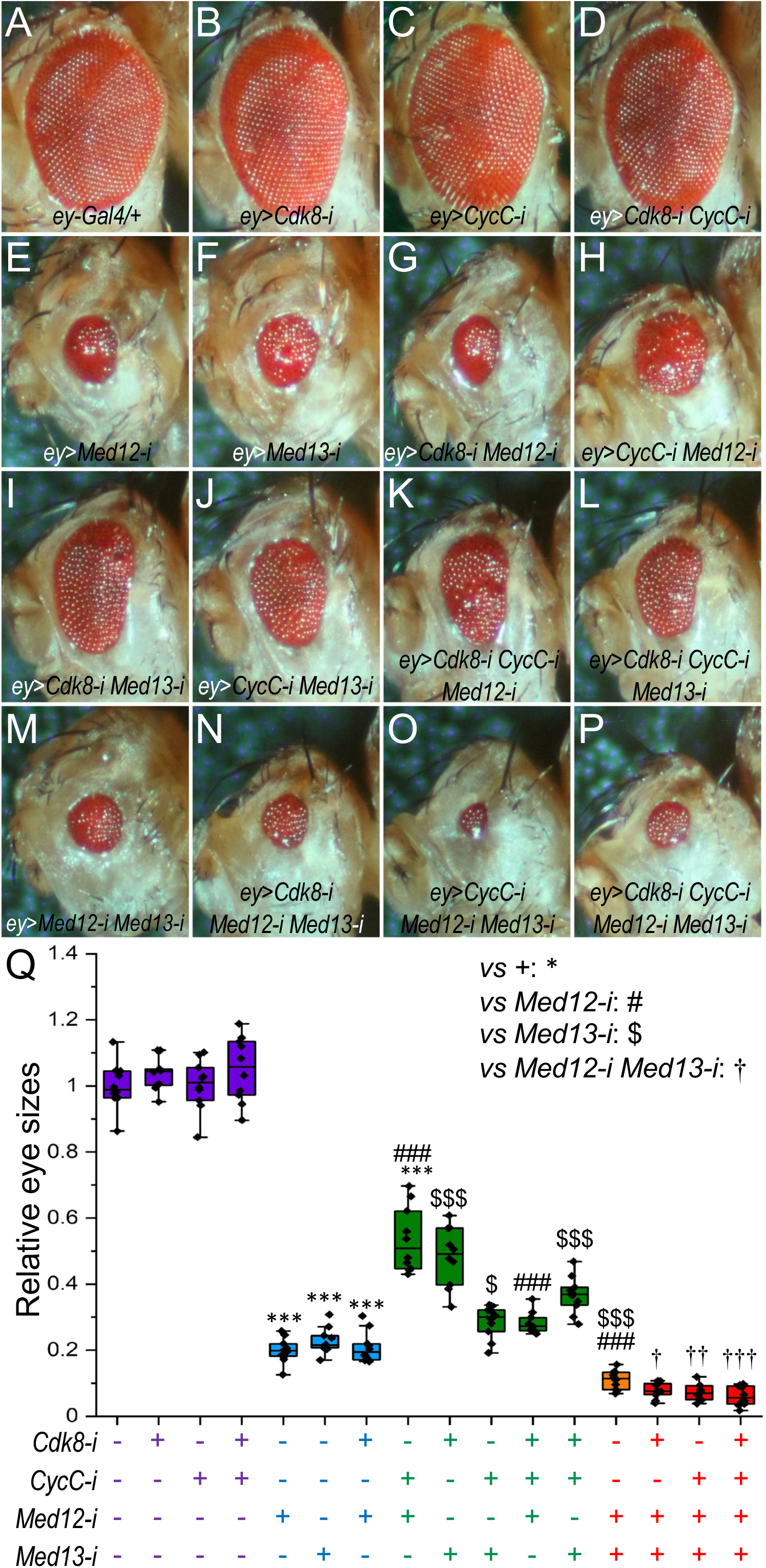
Depletion of various combinations of the CKM subunits led to alterations in eye sizes. Representative adult female eyes were observed for the following genotypes: (A) *ey-Gal4/+* (control); (B) *ey-Gal4/+; UAS-Cdk8-RNAi/+*; (C) *ey-Gal4/+; UAS-CycC-RNAi/+*; (D) *ey-Gal4/+; UAS-Cdk8-RNAi CycC-RNAi/+*; (E) *ey-Gal4/+; UAS-Med12-RNAi/+*; (F) *ey-Gal4/+; UAS-Med13-RNAi/+*; (G) *ey-Gal4/+; UAS-Cdk8-RNAi Med12-RNAi/+*; (H) *ey-Gal4/+; UAS-CycC-RNAi Med12-RNAi/+*; (I) *ey-Gal4/+; UAS-Cdk8-RNAi Med13-RNAi/+*; (J) *ey-Gal4/+; UAS-CycC-RNAi Med13-RNAi/+*; (K) *ey-Gal4/+; UAS-Cdk8-RNAi CycC-RNAi Med12-RNAi/+*; (L) *ey-Gal4/+; UAS-Cdk8-RNAi CycC-RNAi Med13-RNAi/+*; (M) *ey-Gal4/+; UAS-Med12-RNAi Med13-RNAi/+*; (N) *ey-Gal4/+; UAS-Cdk8-RNAi Med12-RNAi Med13-RNAi/+*; (O) *ey-Gal4/+; UAS-CycC-RNAi Med12-RNAi Med13-RNAi/+*; and (P) *ey-Gal4/+; UAS-Cdk8-RNAi CycC-RNAi Med12-RNAi Med13-RNAi/+*. (Q) Eye sizes were quantified with N>10 for each genotype. Significance levels: */#/$/& p<0.05; **/##/$$/&& p<0.01; ***/###/$$$/&&& p<0.001, all based on one-tailed unpaired *t*-tests.

Depleting both Med12 and Med13 resulted in even smaller eyes than depleting either Med12 or Med13 alone, indicating that the hypomorphic and modifiable nature of these phenotypes (Fig. 1M). Notably, depleting CycC-Med12 (Fig. 1H) resulted in larger eyes than depleting Med12 alone (Fig. 1E), while depleting CDK8-Med13 or CycC-Med13 (Fig. 1I and 1J) led to larger eyes than depleting Med13 alone (Fig. 1F), as quantified in Fig. 1Q. Similarly, simultaneous depletion of CDK8 and CycC together with Med12 or Med13 (Fig. 1K, 1L) led to larger eyes than knocking down Med12 or Med13 alone (Fig. 1E, 1F), although the eye size of these flies remained smaller than that of the control. These results sugest a partial rescue of the Med12-RNAi or Med13-RNAi phenotype by depleting CDK8-CycC in general, even though depleting CDK8-Med12 (Fig. 1G) resulted in a similar eye size to depleting Med12 alone (Fig. 1H). The partial rescue implies an antagonistic relationship between Med12-Med13 and CDK8-CycC during *Drosophila* eye development. Furthermore, knocking down CDK8 or CycC in addition to Med12-Med13 together, or knocking down all four CKM subunits (Fig. 1N, 1O, and 1P), resulted in more sever effects than those caused by depleting Med12 and Med13 together (Fig. 1M), as quantified in Fig. 1Q. This indicates that the rescuing effects of CDK8-CycC depend on the residual presence of Med12-Med13. These observations validate the essential roles of the CKM, particularly Med12 and Med13, in eye development.

The quantification of the eye phenotypes using this set of transgenic RNAi lines highlights five distinct classes of RNAi effects (Fig. 1Q): depletion of CDK8, CycC, or both CDK8 and CycC (Fig. 1B, 1C and 1D) has no effects on eye sizes (designated as the Class I). Depleting Med12 or Med13 alone (Fig. 1D and 1E) results in reduced eye sizes (designated as the Class II). Simultaneous depletion of one or two subunits of CDK8-CycC pair partially rescues the small eye phenotype caused by depleting Med12 or Med13 alone (Fig. 1H, 1I, 1J, 1K and 1L, designated as the Class III). Knocking down both Med12 and Med13 (Fig. 1M) leads to smaller eye (designated as the Class IV). Lastly, depleting one or two subunits of CDK8-CycC pair in additional to Med12 and Med13 together (Fig. 1N, 1O, and 1P) results in the most severe small eye phenotypes compared to other combinations (designated as the Class V). These analyses suggest an unexpected antagonistic function between the CDK8-CycC pair and the Med12-Med13 pair *in vivo*, which is intriguing, given that the four subunits of the CKM can be biochemically purified as a complex and are typically assumed to function together as a unit.

We conducted additional experiments to examine the effects of depleting different subunits of the CKM in *Drosophila* wings using the *apterous-Gal4* (*ap-Gal4*) driver, which is specifically expressed in the dorsal compartment of wing discs (37). To simplify our subsequent analyses, we focus on the following representative lines for each class: *UAS-Cdk8-i, CycC-i* for Class I, *UAS-Med12-i* for Class II, *UAS-Cdk8-i, CycC-i, Med12-i* for Class III, *UAS-Med12-i, Med13-i* for class IV, and *UAS-Cdk8-i, CycC-i, Med12-i, Med13-i* for Class V. When we depleted CDK8, the adult wings exhibited a characteristic downward curl towards the ventral side (Fig. S1B), consisting with the known inhibitory effects of CDK8 on cell proliferation (11, 38). In contrast, depleting Med12 alone (Fig. S1C) led to severe disruption of wing morphology, accompanied by the formation of blisters and the shrinking of wing blades. Similar effects were observed when CDK8 and CycC was co-depleted along with Med12 (Fig. S1D). Depleting both Med12 and Med13 simultaneously, or all four subunits together, led to pupal lethality. Animals dissected from the pupal case exhibited a severe disruption in wing morphology (Fig. S1E and S1F). Due to difficulties in reliably quantifying these wing phenotypes, our subsequent analyses at the cellular level were focused on wing discs.

### Distinct roles of the CKM subunits in regulating cell proliferation in wing discs

Alterations in organ sizes at the cellular level can result from changes in cell proliferation, apoptosis, or a combination of both. To investigate whether the reduction in organ size caused by depleting Med12-Med13 is due to apoptosis, we immunostained wing discs using an anti-cleaved Dcp-1 (death caspase-1) antibody to identify apoptotic cells (39). In control discs (Fig. S2A), cells positive for cleaved Dcp-1 could occasionally be observed in the wing pouch area. When we used *ap-Gal4* to deplete Med12-Med13 together or all four CKM subunits simultaneously, there was no obvious alteration in the distribution or the number of Dcp-1 positive cells (Fig. S2B and S2C). This suggests that the reduction of Med12-Med13 or the entire CKM has little effect on apoptosis.

Next, we investigated the effect of depleting the CKM subunits on cell proliferation in wing discs using an antibody recognizing phosphorylated Histone H3 at Serine 10 (PH3), which marks mitotic cells. In control wing discs, PH3-positive cells are randomly distributed throughout the wing pouch area, and the ratio of mitotic cells in the dorsal compartment to those in the ventral compartment is approximately equal to one (Fig. 2A; quantified in Fig 2G). Depletion of both CDK8 and CycC using *ap-Gal4* led to a significant increase in the number of PH3-positive cells in the dorsal compartment compared to the ventral compartment in the same wing discs (Fig. 2B; Fig 2G). In contrast, depleting Med12 alone (Fig. 2C) or both Med12 and Med13 (Fig. 2E) did not affect the number of PH3-positive cells between the dorsal and ventral compartments of the same discs (Fig. 2G), suggesting that the reduction of Med12-Med13 does not affect mitosis. Notably, when CDK8 and CycC were depleted in addition to Med12, a significant increase in the number of PH3-positive cells in the dorsal compartment was observed (Fig. 2D; Fig 2G). Similar observations were made when Med12 and Med13 were depleted alongside CDK8 and CycC (Fig. 2F; Fig 2G). These findings suggest that CDK8-CycC, but not Med12-Med13, negatively regulates cell proliferation.

**Fig. 2.**
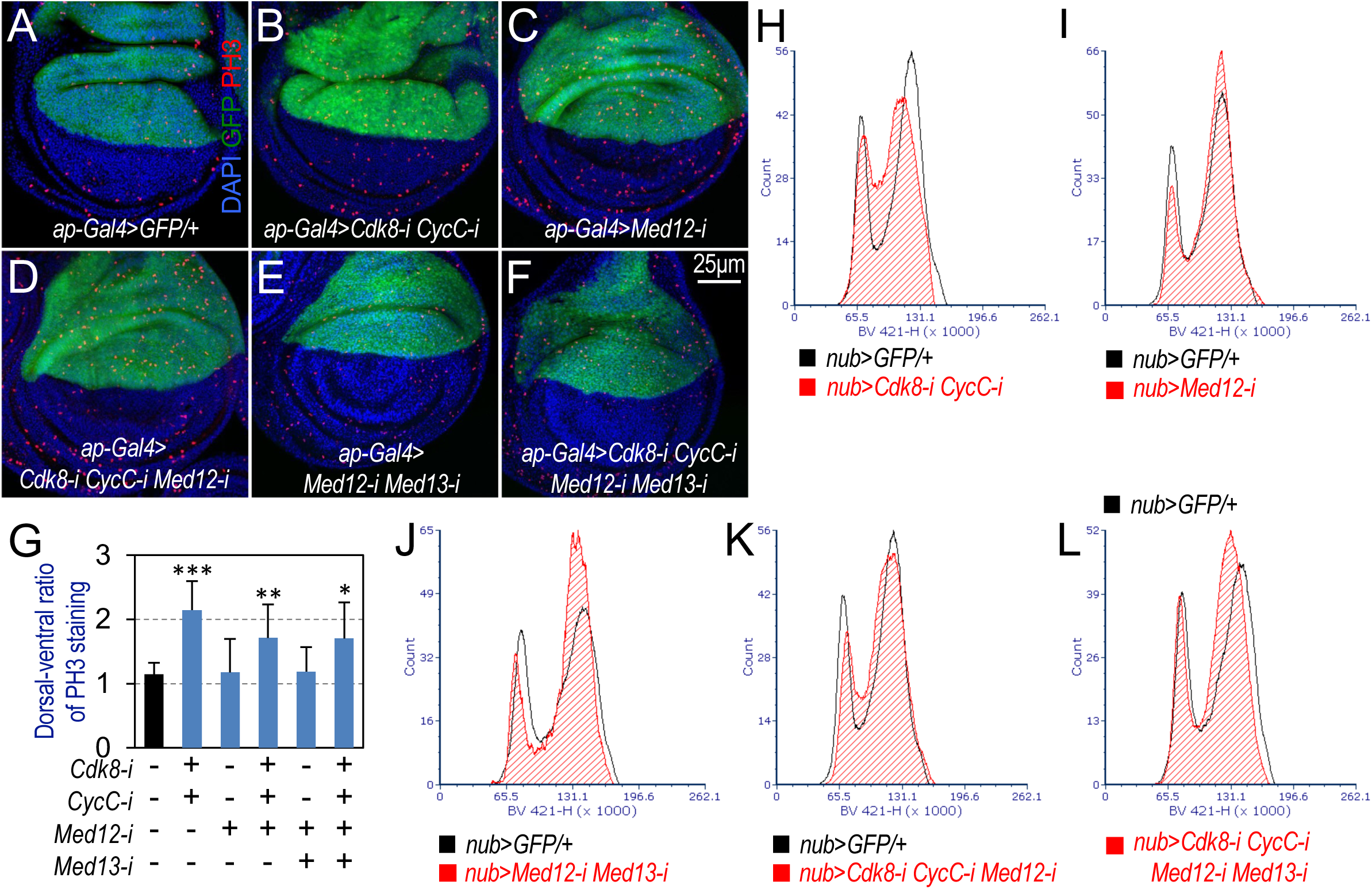
Distinct effects of CKM subunit depletion on cell-cycle progression. Representative confocal images of the wing pouch area from wing imaginal discs of third instar larvae at the wandering stage are presented. Wing discs were stained with DAPI (blue) and anti-PH3 antibody (red), while GFP (green) labeled the dorsal compartment. The specific genotypes analyzed are as follows: (A) *ap-Gal4>UAS-2XGFP/+*; (B) *ap-Gal4>UAS-2XGFP/+; UAS-Cdk8-RNAi CycC-RNAi/+*; (C) *ap-Gal4>UAS-2XGFP/+; UAS-Med12-RNAi/+*; (D) *ap-Gal>UAS-2XGFP/+; UAS-Cdk8-RNAi CycC-RNAi Med12-RNAi/+*; (E) *ap-Gal4>UAS-2XGFP/+; UAS-Med12-RNAi Med13-RNAi/+*; and (F) *ap-Gal4>UAS-2XGFP/+; UAS-Cdk8-RNAi CycC-RNAi Med12-RNAi Med13-RNAi/+*. Scale bar in panel F: 25μm. (G) Quantification of the PH3 signal number was performed using five wing discs of each genotype. * p<0.05; ** p<0.01; *** p<0.001. (H-L) Analyses of CKM subunit depletion effects on cell-cycle progression using flow cytometry. Histogram of flow cytometry counts from DyeCycle violet staining are presented: (H) *nub-Gal4/+; UAS-mCD8::GFP/+* (black) vs. *nub-Gal4/+; UAS-mCD8::GFP/UAS-Cdk8-RNAi CycC-RNAi* (red); (I) *nub-Gal4/+; UAS-mCD8::GFP/+* (black) vs. *nub-Gal4/+; UAS-mCD8::GFP/UAS-Med12-RNAi* (red); (J) *nub-Gal4/+; UAS-mCD8::GFP/+* (black) vs. *nub-Gal4/+; UAS-mCD8::GFP/UAS-Med12-RNAi Med13-RNAi* (red); (K) *nub-Gal4/+; UAS-mCD8::GFP/+* (black) vs. *nub-Gal4/+; UAS-mCD8::GFP/UAS-Cdk8-RNAi CycC-RNAi Med12-RNAi* (red); and (L) *nub-Gal4/+; UAS-mCD8::GFP/+* (black) vs. *nub-Gal4/+; UAS-mCD8::GFP/UAS-Cdk8-RNAi CycC-RNAi Med12-RNAi Med13-RNAi* (red).

To further analyze the effect of depleting the CKM subunits on cell-cycle progression, we used the Fluorescent Ubiquitination-based Cell Cycle Indicator (FUCCI) system in *Drosophila*, known as Fly-FUCCI. This system offers a sophisticated *in vivo* assessment of cell-cycle status in complex developmental contexts (40). Using two fluorescent reporters, GFP-dE2f1_1-230_ and mRFP-dCycB_1-266,_ simultaneously expressed in all cells of developing tissues, we can distinguish cell-cycle phases: G1 phase emits green fluorescence, S phase emits red, and G2 phase appears yellow due to the co-expression of both probes (40). Except for the Zone of Nonproliferating Cells (ZNC), EGFP- and RFP-labeled cells are randomly distributed in the remaining regions of the normal wing discs (Fig. S3A), suggesting asynchronous cell division.

The ZNC comprises a stripe of G1 arrested cells in the posterior part of the DV boundary (marked by green cells within the white dotted rectangle in Fig. S3A; quantified in Fig. S3G), along with a stripe of G1 arrested cells along the center of the DV boundary; this central stripe is flanked by two stripes of G2 arrested cells in the anterior part of the DV boundary (41). Depleting both CDK8 and CycC in the wing pouch region of wing discs using *nub-Gal4* significantly increased the number of red cells (Fig. S3B; Fig. S3G), suggesting that CDK8-CycC plays a negative regulatory role in S phase entry. This observation supports the model suggesting that CDK8 serves as a negative regulator of E2F1, the key transcription factor controlling the G1-S phase transition (11, 12, 42). In contrast, depletion of Med12 alone or Med12 and Med13 together did not significantly affect the number of cells in S phase (Fig. S3C, Fig. S3E; Fig. S3G). However, simultaneous depletion of CDK8, CycC, and Med12 (Fig. S3D; Fig. S3G), or all four CKM subunits (Fig. S3F; Fig. S3G), increases the population of cells in S phase compared to the control (Fig. S3A; Fig. S3G). Together, these observations suggest that depleting CDK8-CycC enhances cell proliferation by promoting entry into both S phase and mitosis, whereas Med12-Med13 depletion lacks this effect.

In addition to its effects on cell proliferation, we observed distinct alterations in the patterning of the FUCCI signal within the ZNC resulting from the depletion of various combinations of the CKM. Depleting both CDK8 and CycC had no discernible effect on the FUCCI patterns in the ZNC (Fig. S3B), as compared to the control (Fig. S3A). In contrast, the four remaining representative combinations of the CKM, where Med12 was depleted either alone or together with other subunits, resulted in the absence of the characteristic FUCCI patterns within the ZNC, as indicated by white dotted rectangles in Fig. S3C-S3F. These differences indicate that depleting Med12 disrupts the patterning or differentiation within the ZNC, while depleting CDK8-CycC does not significantly affect them.

To further investigate the effect of depleting specific combination of CKM subunits on cell-cycle progression, we utilized flow cytometry to analyze the DNA content of cells in the wing disc. Using the *nub-Gal4* driver, we depleted representative CKM subunit combinations and analyzed GFP-labeled cells (*UAS-GFP*). Depletion of both CDK8 and CycC resulted in a notable increase in the population of cells in the S phase, rising from 22.73% in the *nub-Gal4* heterozygous control to 35.97% (Fig. 2H; Fig. S4B; Fig. S4G). In contrast, depleting Med12 alone (Fig. 2I; Fig. S4C), or Med12-Med13 in combination (Fig. 2J; Fig. S4E), did not significantly alter the population of cells in the S phase (Fig. S4G). However, co-depletion of Med12 with CDK8-CycC increased the S phase cell population, reaching 53.82% (Fig. 2K; Fig. S4D; Fig. S4G). Moreover, depletion of all four CKM subunits caused a shift of the G2/M peak towards the G1 peak, indicating an increased transition of cells from the S phase to the G2 phase (Fig. 2L; Fig. S4F; Fig. S4G). These findings are consistent with the results of the FUCCI system, suggesting that depleting CDK8-CycC increases the population of S-phase cells, while depleting Med12-Med13 has little effect on S-phase progression in wing discs.

### Med12 regulates the expression of genes involved in neuronal differentiation

To elucidate the mechanisms underlying the distinct effects of depleting different CKM subunits on cell proliferation and on tissue size, we conducted single-cell RNA sequencing (scRNA-seq) analyses using the 10x Genomics platform. Specifically, we utilized the *ap-Gal4>GFP* system to deplete three representative combinations of CKM subunits, namely CDK8-CycC, Med12 alone, and CDK8-CycC-Med12. We used the *ap-Gal4>GFP* heterozygous wing discs as the control. To identify differences in gene expression among samples, we integrated the scRNA-seq data from four experimental conditions and visualized the cell distributions using the t-Distributed Stochastic Neighbor Embedding (t-SNE) method (43). This approach enabled us to uncover clusters of cells with similar gene expression profiles and compare them across samples with different genetic perturbations. We observed that most cells formed cohesive clusters without clear boundaries. We further characterized these clusters using specific marker genes for distinct subregions of wing disc epithelium cells. Notably, when we highlighted cells with landmark genes such as *nub* (a specific marker for the wing pouch cells, Fig. S5A), *teashirt* (*tsh*) for the notum cells (Fig. S5B), or *pannier* (*pnr*) for cells at the tip of notum (Fig. S5C), the cells clustered together accurately. This indicates precise identification of different cell populations in our samples.

We next applied the t-SNE method to visualize the data from GFP-positive cells in which CDK8-CycC, Med12 alone, or CDK8-CycC-Med12 were depleted. The scRNA-seq data from GFP-positive cells were categorized into 15 distinct clusters, denoted as cluster 0 to cluster 14. Our primary focus was on the clusters that encompassed most epithelial cells and exhibited close clustering, which included clusters 0, 1, 2, 3, 4, 6, 7, 8, 9, and 13 (see Discussion for the “satellite” clusters). Compared to the control (Fig. S6A), depleting CDK8-CycC did not significantly alter the distribution of cells within these clusters (Fig. S6B). However, depleting Med12 led to a substantial reduction in the cell population within cluster 13, decreasing from 1.41% to 0.38% (Fig. S6C and S6E). Likewise, the simultaneous depletion of CDK8, CycC, and Med12 resulted in a significant decrease in the cell population of cluster 13, from 1.41% to 0.13% (Fig. S6C and S6E). These results indicate that Med12 plays a distinct role from CDK8-CycC in the regulation of gene expression within cluster 13 cells.

To identify the cluster 13 cells, we initially identified the genes with high expression levels within this cluster. Subsequently, we conducted a Gene Ontology (GO) analysis using DAVID GO (Database for Annotation, Visualization, and Integrated Discovery Gene Ontology (44)) on these genes. The GO analysis revealed that the enriched genes within cluster 13 cells were closely associated with neuronal cell differentiation, including categories such as ‘Neurogenesis’ and ‘Notch signaling pathway’ (Fig. 3A). This finding is consistent with the observations derived from the FUCCI system at ZNC (Fig. S3), where cells develop into adult wing margin neuronal cell (41). Collectively, these results suggest that Med12-Med13 plays a more significant role than CDK8-CycC in regulating the expression of genes involved in neuronal differentiation.

**Fig. 3.**
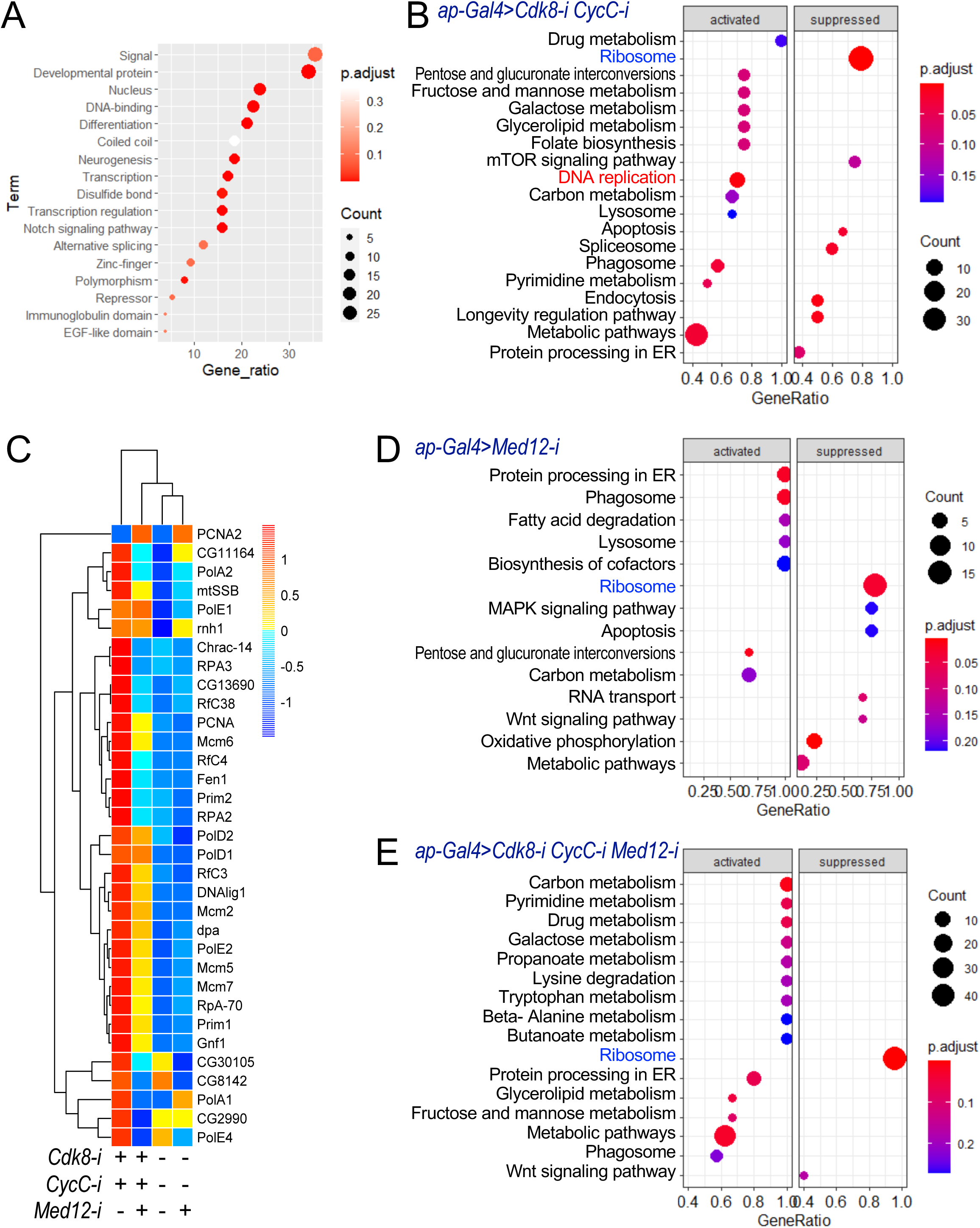
scRNA-seq data from wing discs after depleting different CKM subunits. (A) A Dotplot illustrating DAVID Gen Oncology (GO) analysis results for cluster 13. (B/D/E) Dotplots showing KEGG pathway analyses comparing differentially expressed genes between GFP+ cells to GFP-cells in Nub+ population of the following genotypes: (B) *ap-Gal4>UAS-2XGFP/+; UAS-Cdk8-RNAi CycC-RNAi/+*; (D) *ap-Gal4>UAS-2XGFP/+; UAS-Med12-RNAi/+*; and (E) *ap-Gal>UAS-2XGFP/+; UAS-Cdk8-RNAi CycC-RNAi Med12-RNAi/+.* (C) A heatmap depicting relative gene expression changes in the ‘DNA replication’ category across the same genotypes as in panel B, D, and E. The genotype for the control: *ap-Gal4>UAS-2XGFP/+; +*.

### Distinct impacts on the transcriptome caused by depletion of different CKM subunits

One caveat of this analysis is that *ap-Gal4* expression is confined to the dorsal compartment of the wing discs, which harbors more sensory organ precursors compared to the ventral compartment (37, 45). These sensory organ precursors eventually differentiate into different sensory organs on the notum of adult flies, including bristles, campaniform sensilla, and chordotonal organs. To mitigate potential bias resulting from dorsal specific expression of *ap-Gal4* in wing discs, we reanalyzed the scRNA-seq data, with particular attention to the transcriptome of *nubbin* (*nub*)-positive cells. *Nub* is expressed symmetrically along the dorsoventral (DV) boundary within the wing pouch area (46). Specifically, we analyzed gene expression profiles by comparing cells located in the dorsal compartment (positive for both *nub* and GFP expression) with those in the ventral compartment (positive for *nub* but lacking GFP expression).

To identify differentially expressed genes caused by depleting different CKM subunits, we excluded genes that did not exhibit symmetrical expression along the DV boundary in control sample. As indicated by the Venn diagrams (Fig. S7A), we excluded 2,611 genes that showed significantly higher expression levels in the dorsal compartment of both the control sample and the *ap-Gal4>UAS-Cdk8-i CycC-i* sample. Therefore, we focused our analysis on the 2,280 genes that were uniquely upregulated in cells with CDK8 and CycC depletion (Fig. S7A). Additionally, we conducted further analyses on 1,733 genes that were upregulated in cells with Med12 depletion (Fig S7B) and 2,097 genes that exhibited increased expression in cells with simultaneous depletion of CDK8, CycC, and Med12 (Fig. S7C, as shown below). Furthermore, we also examined 1,013 gene that were down-regulated by depleting CDK8 and CycC (Fig. S7D), 1,511 genes that were reduced by depleting Med12 (Fig. S7E), and 554 genes that were down-regulated by simultaneous depletion of all three subunits (Fig. S7F).

To identify the significant and uniquely affected genes in each sample, we conducted pathway enrichment analyses using the KEGG (Kyoto Encyclopedia of Genes and Genomes) database. Our analyses revealed that genes affected by the depletion of CDK8 and CycC were enriched in pathways such as the ‘DNA replication,’ ‘Ribosome,’ and several metabolic pathways (Fig. 3B). To visualize changes in the genes involved in the ‘DNA replication’ pathway, we generated a heatmap that indicated upregulation (Fig. 3C). These findings align with our earlier observations indicating an increase in the S-phase cell population and cell proliferation (Fig. 2), further supporting our previous report that CDK8 negatively regulates E2F1 (11). However, when Med12 was depleted alone, the ‘DNA replication’ pathway did not show enrichment in this analysis (Fig. 3D), indicating that Med12 depletion did not notably affect DNA replication. Instead, our analyses revealed significant alterations in the ‘MAPK signaling pathway,’ ‘Wnt signaling pathway,’ ‘Oxidative phosphorylation,’ and ‘RNA transport’ pathways upon Med12 depletion (Fig. 3D). Although the genes affected by depleting CDK8, CycC, and Med12 were not enriched in the ‘DNA replication’ pathway (Fig. 3E), our heatmap showed that major E2F1 target genes, such as *PCNA*, *MCM2*, *MCM5* and *MCM6*, were upregulated in both *ap-Gal4>UAS-Cdk8-i CycC-i* and *ap-Gal4>UAS-Cdk8-i CycC-i Med12-i* samples, but not in *ap-Gal4/+* or *ap-Gal4>UAS-Med12-i* samples (Fig. 3C). These results reveal a distinct function of CDK8-CycC in regulating the expression of genes involved in the ‘DNA replication’ pathway, supporting a negative regulation of E2F1 transcriptional activity of E2F1 by CDK8-CycC, while the impact of Med12 depletion appears to be marginal. Furthermore, besides the specific roles of CDK8-CycC in DNA replication and Med12 in Wnt signaling, our analysis revealed that depleting different CKM subunits also down-regulated several common pathways, including the ‘Ribosome’ and ‘Metabolic pathways’ (Fig. 3B, Fig. 3D, and Fig. 3E).

The data above suggest that depleting CDK8-CycC or Med12 has distinct impacts on different pathways, with overlapping effects on specific pathways. Furthermore, depleting CDK8, CycC, and Med12 led to an additive effect, combining the individual effects of depleting CDK8-CycC or Med12 alone.

### Effects of the CKM subunits on E2F1-dependent transcription

Given the potential influence of sequencing depth, library preparation biases, and other factors on the sensitivity and reliability of scRNA-seq analysis (47), we sought to further verify the effect of CKM subunit depletion on E2F1 target gene expression using the Hybridization Chain Reaction Fluorescence In Situ Hybridization (HCR RNA-FISH) assay. This method, known for its high specificity, multiplexing capability, and single-molecular resolution, enables quantitative analysis of RNA expression at the single-cell level (48). Specifically, we focused on three conserved E2F1 target genes: *CycE* (*Cyclin E*), *mcm5* (*minichromosome maintenance 5*), and *stg* (*string*; encoding the *Drosophila* CDC25 phosphatase homolog) (11, 49–51). The expression of these genes, particular evident for *Mcm5* (Fig. 4A’), in the wing disc displays a characteristic expression pattern in the ZNC that correlates with the cell-cycle distribution in this region (41). As expected, depletion of Dp in the posterior compartment reduced the expression of *CycE* (Fig. 4B), *Mcm5* (Fig. 4B’), and *stg* (Fig. S8D; quantified in Fig. 4G), compared to cells in the anterior compartment of the same discs. Conversely, depletion of *Rbf1* resulted in increased expression of *CycE* (Fig. 4C), *Mcm5* (Fig. 4C’), and *stg* (Fig. S8E). Moreover, overexpression of *CycE* (Fig. 4D) increased the expression of *Mcm5* (Fig. 4D’) but had no obvious effects on *stg* levels (Fig. S8F).

**Fig. 4.**
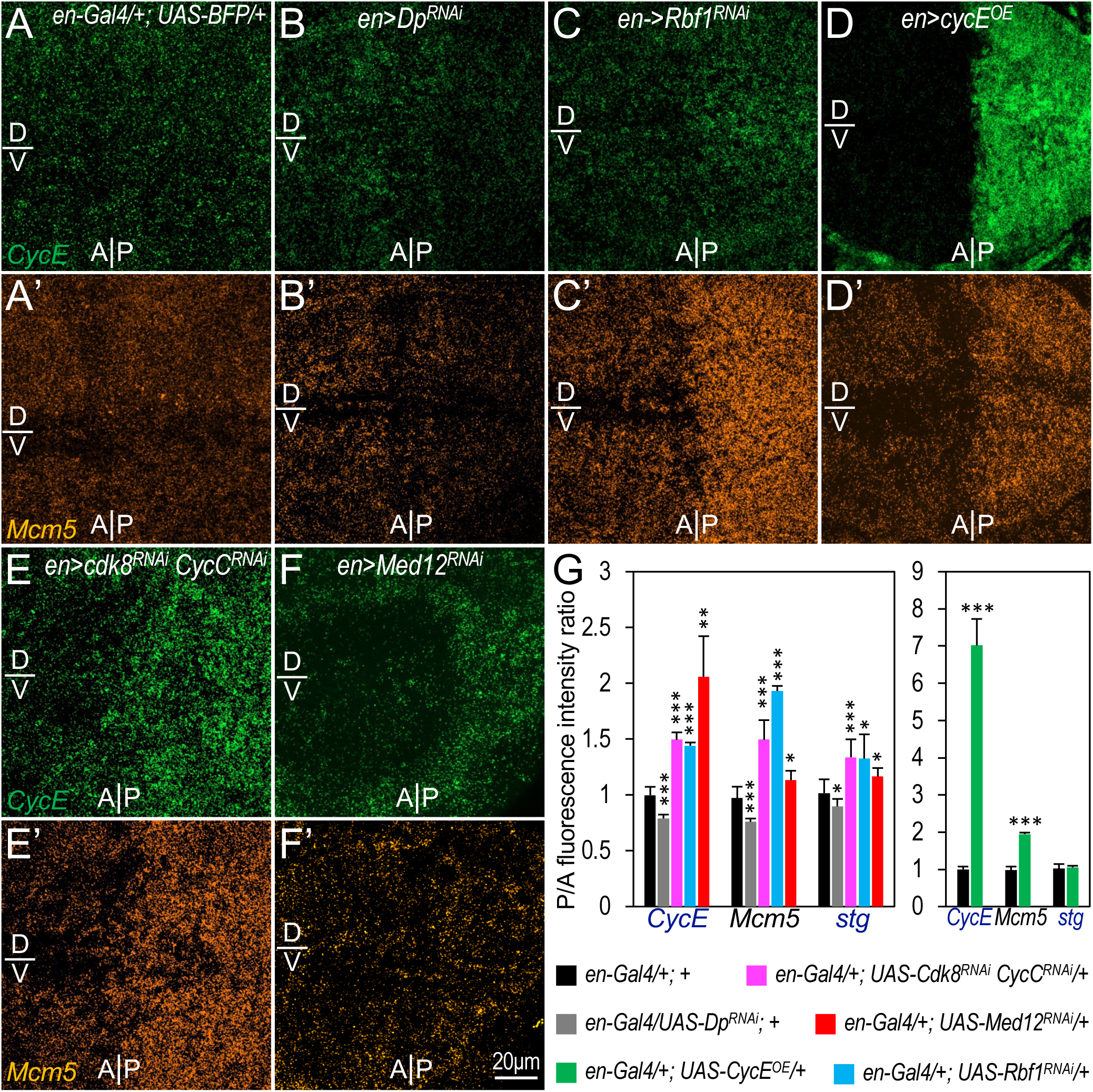
The role of CKM in regulating the expression of E2F1 target genes. Representative confocal images display mRNA transcripts of *CycE* (green) and *Mcm5* (orange), detected using the HCR RNA-FISH technique within the wing pouch region of wing discs. Supplementary images illustrating the expression of BFP and *stg* are provided in Suppl. Fig. S8. Genotypes: (A) *en-Gal4/+; UAS-BFP/+* (control); (B) *en-Gal4/+; UAS-Dp^RNAi^/UAS-BFP*; (C) *en-Gal4/+; UAS-Rbf1^RNAi^/UAS-BFP*; (D) *en-Gal4/+; UAS-CycE^+^/UAS-BFP*; (E) *en-Gal4/+; UAS-Cdk8-RNAi CycC-RNAi/UAS-BFP*; and (F) *en-Gal4/+; UAS-Med12-RNAi/UAS-BFP*. The scale bar in panel F’: 20 μm. (G) Quantification of *CycE*, *Mcm5*, and *stg* expression reveals relative expression levels of these genes in the dorsal-posterior compartment, normalized to the dorsal-anterior compartment of the same disc. ‘A/P’ denotes the anterior-posterior compartment boundary, while ‘D/V’ indicating the dorsal-ventral boundary. Three to five discs for each genotype, color-coded below, were analyzed. * p<0.05, ** p<0.01, *** p<0.001 (one-tailed unpaired *t*-tests).

Importantly, depletion of Cdk8 and CycC in the posterior compartment cells markedly increased the expression of these E2F1 target genes compared to those in the anterior compartment (Fig. 4E/E’, and Fig. S8B/B’; quantified in Fig. 4G). This observation supports the notion that CDK8-CycC negatively regulates E2F1-dependent transcription (11). Similarly, depletion of Med12 in the posterior compartment cells also increased the expression of these E2F1 target genes (Fig. 4F/F’, and Fig. S8C/C’; quantified in Fig. 4G). Notably, mRNA transcript detection via the HRC assay may offer a more sensitive and direct measure of E2F1-dependent transcription compared to the scRNA-seq analysis. These findings suggest the involvement of CKM, rather than a CKM-independent function of CDK8-CycC, in regulating E2F1-dependent transcription.

### The role of CKM subunits in regulating ribosomal protein transcription

To further explore the mechanism underlying the phenotypic difference between CDK8-CycC depletion and Med12-Med13 depletion, we directed our attention to KEGG pathway analysis, revealing notable changes in a substantial number of genes linked to ‘Ribosome’ pathway (Fig. 3B, Fig. 3D, Fig. 3E). This is intriguing due to the essential role of ribosomes in protein synthesis and cellular growth.

In both *Drosophila* and humans, the mature 80S ribosomes comprise a small 40S module with 18S ribosomal RNA (rRNA) and 33 ribosomal proteins, and a large 60S module containing 28S, 5.8S, and 5S rRNAs, along with 47 ribosomal proteins (52, 53). As shown in the heatmap (Fig. 5A), 74 out of 80 genes encoding ribosomal proteins in the ‘Ribosome’ pathway were downregulated in the Med12-RNAi sample. Among these, 72 genes showed consistent downregulation in samples with CDK8 and CycC depletion. Six ribosomal genes – *tko*, *bonsai*, *RpL34a*, *RpLS10a*, *RpS15Ab*, and *RpS19b* – are upregulated in Med12-depleted samples (Fig. 5A). Except for *bonsai*, the depletion of CDK8 and CycC had no obvious effect on the expression of the other five ribosomal genes (Fig. 5A). Notably, *tko* and *bonsai* encode mitochondrial ribosomal protein S12 and S15, respectively. These findings suggest that CKM may modulate the expression of most ribosomal genes, with minimal difference observed among CKM subunits.

**Fig. 5.**
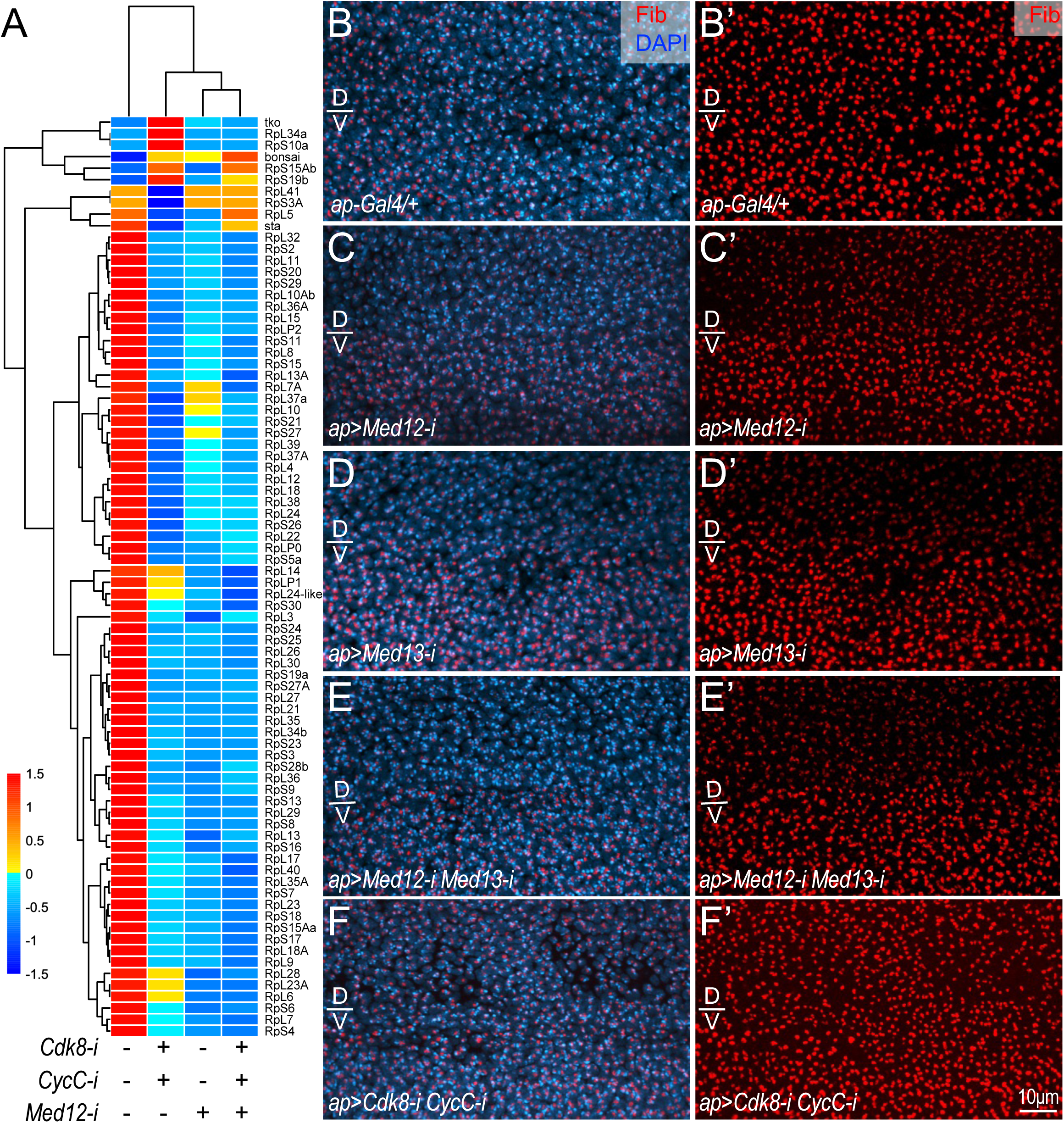
The distinct effects of depleting CKM subunits on the transcription of ribosomal genes and fibrillarin expression. (A) Heatmap displaying gene changes in the Ribosome category for the following genotypes: ‘*ap-Gal4>UAS-2XGFP/+*’; ‘*ap-Gal4>UAS-2XGFP/+; UAS-Cdk8-RNAi CycC-RNAi/+*’; ‘*ap-Gal4>UAS-2XGFP/+; UAS-Med12-RNAi/+*’, and ‘*ap-Gal>UAS-2XGFP/+; UAS-Cdk8-RNAi CycC-RNAi Med12-RNAi/+*’. (B-F) Representative confocal images focused on the wing pouch region of wing discs with anti-fibrillarin (Fib; red) and DAPI (blue) staining for the following genotypes: (B) *ap-Gal4/+*; (C) *ap-Gal4; UAS-Med12-RNAi/+*; (D) *ap-Gal4; UAS-Med13-RNAi/+*; (E) *ap-Gal4; UAS-Med12-RNAi Med13-RNAi/+*; and (F) *ap-Gal/+; UAS-Cdk8-RNAi CycC-RNAi/+.* C’-F’ display the anti-Fib (red) alone, with ‘D/V’ indicating the dorsal-ventral boundary. The scale bar in panel F’: 10μm.

We next compared the effects of depleting different CKM subunits on fibrillarin, a conserved rRNA 2’-O-methyl transferase crucial for pre-rRNA processing and serving as a marker for the nucleolus (54). The methyltransferase fibrillarin forms a complex with the NOP56-NOP58 heterodimer and the RNA binding protein 15.5K (NHP2L1), as well as ribonucleoprotein (snoRNP) complexes containing box C/D family small nucleolar RNAs (snoRNAs) (55, 56). Guided by snoRNAs, this complex catalyzes post-transcriptional 2’-O-methylation (2’-O-Me) of the ribose in rRNAs within the nucleolus. 2’-O-Me is a prevalent eukaryotic rRNA modifications that stabilizes the secondary and tertiary structures of rRNA, protecting them from hydrolysis (55, 56). In control wing discs (*ap-Gal4* heterozygous), fibrillarin proteins are expressed in all cells within the wing pouch area of wing discs (Fig. 5B, 5B’). Using *ap-Gal4* to deplete *Med12* (Fig. 5C, 5C’), *Med13* (Fig. 5D, 5D’), or both (Fig. 5E, 5E’), significantly reduced fibrillarin protein levels in the dorsal compartments of the wing discs compared to their ventral counterparts. In contrast, depletion of CDK8-CycC did not notably alter the fibrillarin levels when comparing cells in the dorsal compartment to those in the ventral compartment (Fig. 5F, 5F’). These observations suggest that depleting Med12-Med13 may lead to defective pre-rRNA processing and ribosomal biogenesis.

The mechanism underlying the reduced levels of fibrillarin protein upon depletion of Med12-Med13, but not CDK8-CycC, is unknown. In yeast, Nop1 (fibrillarin) interacts with both Nop56 and Nop58; mutation of these proteins led to impaired pre-rRNA methylation and processing, as well as defects in ribosome assembly and growth (57, 58). These interactions and effects on cell growth are conserved in mammalian cells (54). It is thus conceivable that depleting Med12-Med13, as opposed to CDK8-CycC, may result in a more pronounced disruption of ribosome biogenesis, protein translation, and cellular growth, mediated by fibrillarin.

### Asymmetric interdependency among the CKM subunits

Previously, we observed the destabilization of CycC protein in *cdk8* mutant larvae, whereas the stability of CDK8 protein remained unaffected in *cycC* mutants (29). Thus, we asked whether the loss of one subunit of the CKM could affect the levels of other three CKM subunits. Addressing this question is challenging due to the lack of antibodies specifically targeting the CKM subunits that are suitable for cell biological and biochemical assays. To overcome this obstacle, we used the CRISPR-Cas9 technique to introduce EGFP (enhanced green fluorescent protein) tags into the endogenous loci of *Cdk8*, *Med12*, and *Med13*. In addition, we added an RFP (red fluorescent protein) tag to the endogenous locus of *CycC* (see Materials and Methods for details). Based on structural studies of CKM (24, 59, 60), we positioned these tags at the C-termini of the CKM subunits, designated as CDK8-EGFP, CycC-RFP, Med12-EGFP, and Med13-EGFP. Homozygotes of all EGFP-tagged lines are fully viable and fertile, suggesting that the EGFP tag do not interfere with the normal function of these CKM subunits *in vivo*. However, *CycC-RFP* homozygotes are lethal, indicating that the RFP tag disrupts its normal functions. Nonetheless, the specificity of these lines was verified through PCR and, importantly, through the genetic and cell biological analyses described below.

In wing discs from third instar larvae, CDK8-EGFP (Fig. 6A), CycC-RFP (Fig. 6F), Med12-EGFP (Fig. 6K), and Med13-EGFP (Fig. 6P) are uniformly expressed in the nuclei of all cells. Depletion of CDK8 in the dorsal compartment of the wing disc using *ap-Gal4* significantly reduced CDK8-EGFP level (Fig. 6B). Similarly, depleting CycC reduced CycC-RFP level (Fig. 6H), knocking down Med12 diminished Med12-EGFP levels (Fig. 6N), and depleting Med13 ablated Med13-EGFP levels (Fig. 6T). These expected results validate the specificity and efficacy of both the transgenic RNAi lines targeting the CKM subunits and the EGFP- or RFP-tagged lines for these CKM subunits.

**Fig. 6.**
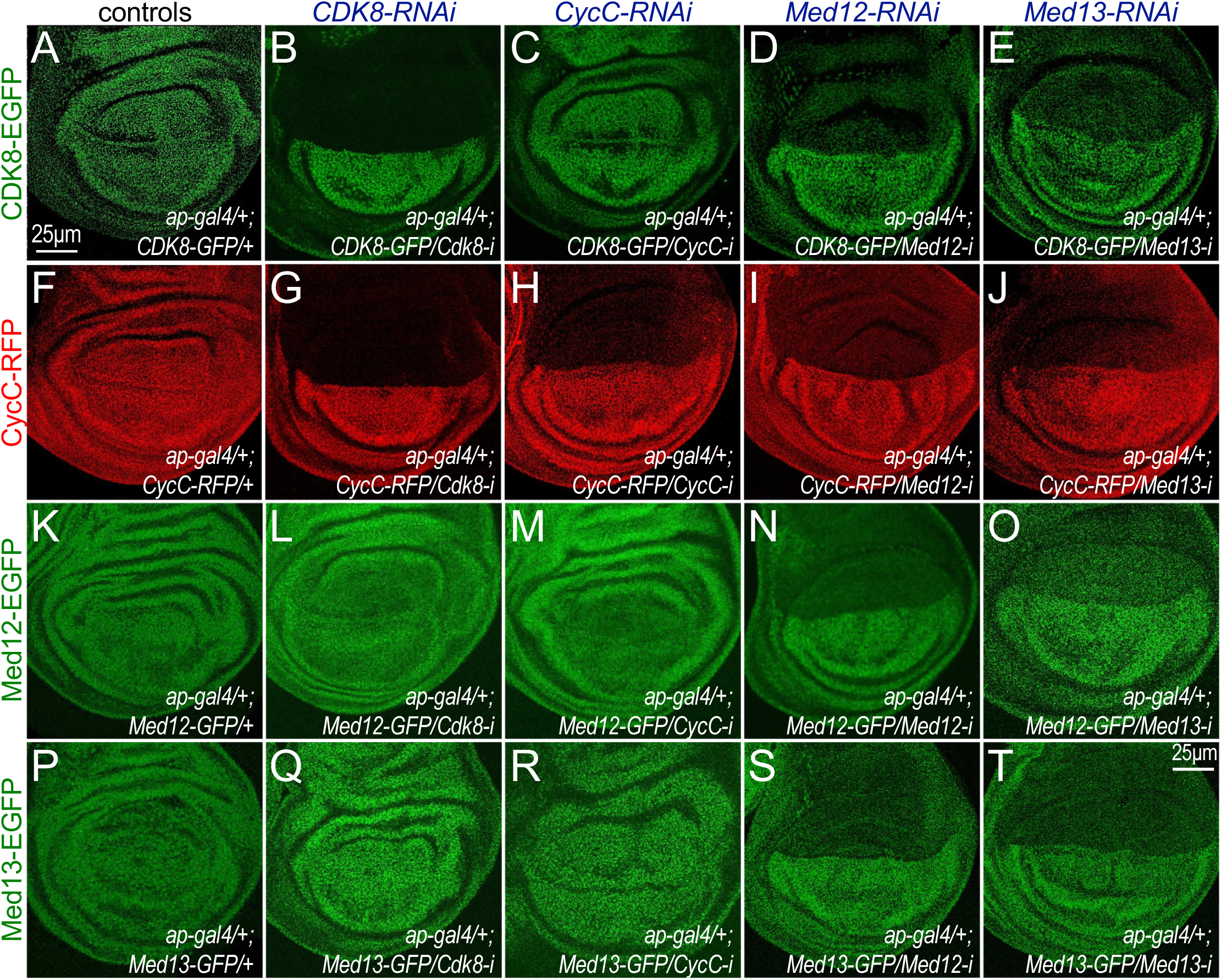
Asymmetric interdependency among CKM subunits. (A-E) Effects of depleting CKM subunits on CDK8-EGFP levels. Genotypes: (A) *ap-Gal4/+; CDK8-EGFP/+*; (B) *ap-Gal4/+; CDK8-EGFP/UAS-Cdk8-RNAi*; (C) *ap-Gal4/+; CDK8-EGFP/UAS-CycC-RNAi*; (D) *ap-Gal4/+; CDK8-EGFP/UAS-Med12-RNAi*; and (E) *ap-Gal4/+; CDK8-EGFP/UAS-Med13-RNAi*. (F-J) Effects of depleting CKM subunits on CycC-RFP levels. Genotypes: (F) *ap-Gal4/+; CycC-RFP/+*; (G) *ap-Gal4/+; CycC-RFP/UAS-Cdk8-RNAi*; (H) *ap-Gal4/+; CycC-RFP/UAS-CycC-RNAi*; (I) *ap-Gal4/+; CycC-RFP/UAS-Med12-RNAi*; and (J) *ap-Gal4/+; CycC-RFP/UAS-Med13-RNAi*. (K-O) Effects of depleting CKM subunits on Med12-EGFP levels. Genotypes: (K) *ap-Gal4/+; Med12-EGFP/+*; (L) *ap-Gal4/+; Med12-EGFP/UAS-Cdk8-RNAi*; (M) *ap-Gal4/+; MED12-EGFP/UAS-CycC-RNAi*; (N) *ap-Gal4/+; Med12-EGFP/UAS-Med12-RNAi*; and (O) *ap-gal4/+; Med12-EGFP/UAS-Med13-RNAi*. (P-T) Effects of depleting CKM subunits on Med13-EGFP levels. Genotypes: (P) *ap-Gal4/+; Med13-EGFP/+*; (Q) *ap-Gal4/+; Med13-EGFP/UAS-Cdk8-RNAi*; (R) *ap-Gal4/+; MED13-EGFP/UAS-CycC-RNAi*; (S) *ap-Gal4/+; Med13-EGFP/UAS-Med12-RNAi*; and (T) *ap-Gal4/+; Med13-EGFP/UAS-Med13-RNAi*.

Importantly, these genetic tools enabled us to define the effect of losing one subunit on the levels of other CKM subunits *in vivo*. We observed that depletion of Med 12 (Fig. 6D) or Med13 (Fig. 6E) in the dorsal compartment also reduced the levels of CDK8-EGFP, while depleting CycC had no effect on level of CDK8-EGFP (Fig. 6C). Notably, depletion of CDK8 or CycC in the dorsal compartment of wing discs also had no effect on the levels of Med12-EGFP (Fig. 6L and Fig. 6M), or the levels of Med13-EGFP (Fig. 6Q and Fig. 6R), suggesting that the levels of Med12 or Med13 are not dependent on CDK8 or CycC. However, depleting Med13 significantly reduced the levels of Med12-EGFP (Fig. 6O), while depleting Med12 also reduced the Med13-EGFP level (Fig. 6S), suggesting that the levels of Med12 and Med13 are interdependent. These results demonstrate that the stability of CDK8 and CycC is dependent Med12 and Med13; the stability of Med12 and Med13 are interdependent on each other, but independent of CDK8 or CycC; and the stability of CycC is dependent on CDK8, but not vice versa (Fig. 7A). Altogether our results reveal an asymmetric interdependency between CDK8-CycC and Med12-Med13, suggesting a more substantial impact of Med12-Med13 disruption on the entire CKM than the loss of CDK8-CycC.

**Fig. 7.**
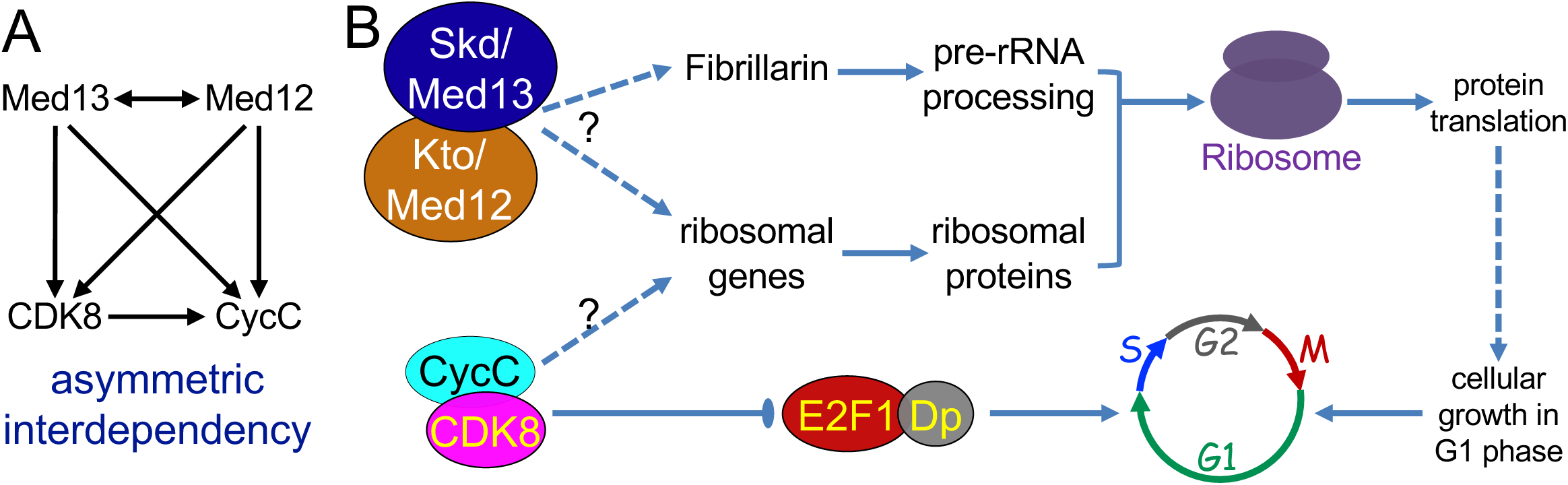
Working models. (A) Dependency diagram summarizing the asymmetric interdependent relationship among CKM subunits. Double-headed arrow between Med12 and Med13 indicates interdependency, while forward arrows denote the direction of dependency. (B) Model: Distinct effects of the CDK8-CycC pair and the Med12-Med13 pair on cellular growth and cell-cycle progression. Loss of all CKM subunits affects ribosomal gene expression. Loss of Med12 or Med13 significantly affects fibrillarin expression, hindering pre-rRNA processing, ribosome maturation, nascent protein translation, and cellular growth. Loss of CDK8 or CycC has minimal effect on fibrillarin expression and pre-rRNA processing sustaining protein translation and cellular growth. This provides the opportunity to manifest the effects of CDK8-CycC on E2F1-dependent transcription and the G1-S phase transition of the cell cycle. Future investigation is needed to understand how CKM subunits modulate the expression of ribosomal genes and fibrillarin.

## Discussion

In this study, we investigated the distinct effects of loss of CKM subunits in *Drosophila* eyes and wings. Depleting Med12 and Med13 leads to severe disruption of cellular growth, likely by interfering with ribosome biogenesis. In contrast, knocking down CDK8 or CycC has a milder effect on cellular growth but affects cell-cycle progression, especially the G1-S phase transition. Our analyses also reveal an asymmetric interdependence in CKM subunit stability: the stability of CDK8 and CycC relies on Med12 and Med13, but not vice versa. These results highlight the different impacts of CKM subunit mutations on cellular growth and proliferation, providing mechanistic insights into their distinct phenotypic consequences *in vivo*.

### Distinct roles of the CKM subunits in regulating *Drosophila* eye and wing development

Our genetic analyses revealed distinct roles for different CKM subunits in eye or wing development. Specifically, CDK8 and CycC are dispensable for proper eye development, as their depletion did not lead to obvious defects in this process. In contrast, Med12 and Med13 are essential for both eye and wing development, as depleting them resulted in severe defects in eye and wing morphology. These phenotypic differences are unlikely due to variations in the efficiency of depleting these four CKM subunits by transgenic RNAi lines, as our observations are consistent with prior reports showing normal eye morphology with *cdk8^K185^* and *cycC^Y5^* mitotic clones, while the presence of *Med12^T241^* mitotic clones in the eye led to severe defects in eye development (27). The efficiencies and specificities of these transgenic lines were confirmed using EGFP-tagged CKM subunits (Fig. 6). The limited impact of CDK8 or CycC depletion on eye size may be attributed to potential underestimation caused by our two-dimensional measurements of the three-dimensional, bulgy shape of adult fly eyes. Therefore, measuring the wing and cell size in wing blade could provide a more accurate assessment. Indeed, depleting CDK8 or CycC using *nub-Gal4* increased the number of wing cells, resulting in a larger wing size, while overexpressing wild-type CDK8 had the opposite effect, reducing wing size (38).

Interestingly, depletion of either Med12 or Med13 resulted in a severe reduction in eye size, partially reversed by additional depletion of CDK8, CycC, or both (Fig. 1Q). Three potential explanations for this apparent antagonistic function between CDK8-CycC and Med12-Med13 can be considered. The first suggests a technical caveat: the expression of multiple shRNAs may impede the efficiency of the RNAi machinery, diminishing the effects of Med12-Med13-RNAi. However, depleting all four CKM subunits resulted in even smaller eyes, indicating that the RNAi machinery is not rate-limiting. The second possibility is that CDK8-CycC may have functions independent of Med12-Med13, indicating CKM-independent roles for CDK8-CycC. This scenario is difficult to reconcile with extensive biochemical and structural studies of the CKM. We favor the third possibility, which suggests that the protein abundance of the four CKM subunits may be differentially affected when the levels of a specific subunit in the complex are diminished (see below).

### Asymmetric interdependency among the four subunits of CKM

In a cryo-electron microscopy study of *Saccharomyces cerevisiae* CKM structure, it was revealed that CDK8 indirectly interacts with Med13 through CycC and Med12 (24). Moreover, Med12 not only extensively interacts with Med13 and CycC, but its N-terminus also directly interacts with CDK8, thereby activating CDK8 alongside CycC (24). Loss of Med13 severely disrupts the interaction between *S. cerevisiae* CKM and the core Mediator complex, while deletion of CDK8 only weakened this interaction (59). The results from our study reveal an asymmetric interdependency among the four CKM subunits, thereby influencing the formation and function of the CKM complex *in vivo*.

Depletion of CDK8 reduces both CDK8 and CycC protein levels, consistent with our previous findings in *cdk8^K185^* mutants during the third instar wandering stage and the white prepupae stage (29). In contrast, depletion of CycC only affects CycC protein levels, with no discernible effect on CDK8 levels, similar to what we observed in *cycC^Y5^* mutant larvae and white prepupae (29). These observations suggest that CycC stability relies on the CDK8, but not vice versa. Therefore, loss of either CDK8 or CycC is expected to disrupt the kinase activity of the CKM. Given that depleting either CDK8 or CycC has little effect on the levels of Med12 and Med13 proteins, the Med12-Med13 dimer may still retain part of the CKM function by interacting with the small Mediator complex. In contrast, depleting either Med12 or Med13 also significantly reduced the levels of CDK8 and CycC proteins and the stability of Med12 and Med13 are mutually interdependent (Fig. 7A). Indeed, loss of either Med12 or Med13 reduced the protein levels of all four CKM subunits. Thus, Med12 or Med13 loss may result in the dissemble of the entire CKM, thereby disrupting the role of CKM in modulating the small Mediator complex and the kinase activity of CKM. This is likely part of the reason for the more severe phenotypic consequences of Med12 or Med13 loss than mutation of CDK8 or CycC.

The asymmetrically interdependence among the four CKM subunits could arise from the corporative stability of protein complexes. This model suggests that the loss of a subunit within a protein complex might expose the intrinsically disordered regions of other subunits, making them vulnerable to degradation by the 20S proteasome (61). Examples for this phenomenon include proteins such as p53 and ornithine decarboxylase (61). Structural analysis reveals that the C-terminus of CDK8 might be shielded by Med12 (24); thus, the loss of Med12 could expose the intrinsically disordered region in the C-terminus of CDK8 to the 20S proteasomes, leading to CDK8 degradation. A similar mechanism may also apply to CycC, Med12, and Med13, which are predicted to contain intrinsically disordered regions (24, 62, 63). This mechanism might complement ubiquitin-dependent 26S proteosome degradation. The E3 ubiquitin ligase for Med13 has been identified as SCF-Fbw7 (64, 65), while the potential E3 ligases for other CKM subunits remain unknown. Further analyses are necessary to fully understand the regulation of this asymmetric interdependency among CKM subunits.

The asymmetric independency among CKM subunits may be more complex in mammalian cells due to the presence of paralogs for three CKM subunits: CDK8 and CDK19, MED12 and MED12L, MED13 and MED13L. Redundancy between these paralogs could contribute to the overall stability of CKM complexes. Notably, the stability of CycC depends on the other three CKM subunits, but not vice versa. It remains uncertain whether this notion explains why CycC lacks paralogs in most animals. Interestingly, CycC does possesses two paralogs in plants and Diplomonadida (6). Whether this asymmetric independency among CKM subunits is conserved in other species remains unknown but may be an intriguing possibility to investigate.

### Distinct effects of depleting CKM subunits on ribosome biogenesis, cellular growth, and cell-cycle progression

Cellular growth is indispensable for cell-cycle progression across various cell types, but the intricate molecular mechanisms bridging these fundamental cellular processes remain not fully elucidated (66). We observed that reducing CDK8 or CycC in wing discs increased the S-phase cell population, and ultimately leading to more but smaller cells in adult wings (38). In contrast, the depletion of Med12 or Med13 had a limited effect on the distribution of cells in different cell-cycle phases.

The 80S ribosomes comprise 80 distinct ribosomal proteins and four types of rRNAs (18S, 28S, 5.8S, and 5S rRNAs), which are derived from pre-rRNAs undergoing modification and processing in the nucleolus, a process facilitated by fibrillarin (53, 54). While depletion of CKM subunits significantly downregulates genes encoding most of ribosomal proteins, depletion of Med12 or both Med12 and Med13 markedly reduces fibrillarin levels. In contrast, depletion of CDK8-CycC does not induce this effect. This discrepancy suggests that loss of CDK8-CycC primarily reduces ribosome abundance, possibly leaving residual ribosomes functional. Conversely, Med12-Med13 depletion may not only diminish ribosome abundance but also disrupt pre-rRNA processing and 80S ribosome assembly through fibrillarin, resulting in more severe defects in ribosome biogenesis, nascent protein translation, and cellular growth (Fig. 7B). Given that cellular growth is a prerequisite for cell division, the severe defects in cellular growth caused by Med12-Med13 depletion may overshadow their influence on E2F1 activities and the G1-S phase transition. In contrast, the depletion or loss of CDK8-CycC results in milder effects on cellular growth and proliferation, enabling the detection of their role in regulating E2F1 activities and the G1-S phase transition (Fig. 7B). This model may explain the distinct effects observed in phenotypes such as those in the *Drosophila* eye and wing (27, 38).

The effect of CDK8-CycC on cell-cycle progression appears to depend on E2F1-regulated transcription (Fig. 7B). Diverse experimental approaches conducted in both *Drosophila* and mammalian cells support the model that CDK8-CycC regulates cell-cycle progression through E2F1-dependent transcription (Fig. 7B). These include the *in vitro* GST-pulldown and co-immunoprecipitation assays, which demonstrate physical interactions between CDK8 and E2F1 (11), as well as *in vitro* CDK8 kinase assays elucidating the phosphorylation of E2F1 as a consequence of this interaction (11, 12). These findings are further complemented by functional and phenotypic analyses, including genetic interactions between CDK8 and E2F1, as well as the impact of CDK8 or CycC loss on the expression of E2F1 target genes (11, 67) (and this work). Considering the established roles of CDK8-CycC within the CKM and the Mediator complex, it was presumed that both the Mediator complex and the CKM coordinate E2F1-dependent transcription (42). In this study, we observed that depleting Med12 in wing disc cells also lead to upregulation of E2F1 target genes such as *CycE*, *mcm5*, and *stg*. This observation further bolsters the notion of the CKM’s involvement in regulating E2F1-dependent transcription.

The mechanism by which CKM modulates the expression of fibrillarin and ribosomal proteins is still unknown. In addition to its role in regulating rRNA stability and processing, fibrillarin can influence rRNA transcription by methylating histone H2A (68, 69). Thus, decreased rRNA levels could potentially result from reduced fibrillarin levels. Nevertheless, whether and how Med12-Med13 regulates the transcription of the *fibrillarin/FBL* gene, and whether this regulation is conserved in mammalian cells, remain unclear. The transcription of *FBL* gene, considered an oncogene with elevated levels in multiple human cancers (54), is directly inhibited by p53 in mammalian cells (70). Interestingly, the C-terminus and the activation domain of p53 directly interact with different Mediator subunits, inducing structural shifts in Mediator and consequently activating the transcription of p53 target genes (71). Moreover, CDK8 has been shown to positively regulated p53-dependent transcription of *p21* in mammalian cells treated with Nutlin3 (14). Furthermore, the oncoprotein MYC has been demonstrated to directly binds to the 5’ upstream region of the *FBL* gene and enhance *FBL* transcription (72). Along these lines, it is established that rDNA transcription relies on RNA polymerase I, with Myc playing a critical role in this process, both in mammals (73) and in *Drosophila* as well (74). Therefore, investigating the regulation of *FBL* transcription by p53 and Myc, as well as the potential role of CKM in this process, in future research will be intriguing.

Notably, CDK8 amplification or up-regulation is observed in various cancer types, including colorectal cancer and melanoma, while its regulatory partner CycC is highly expressed in colorectal-adenocarcinoma, leukemia, and lymphoma cells (33). Additionally, mutations in Med12 in exon 1 and exon 2 are prevalent in over 60% of uterine leiomyomas across different countries (75). Furthermore, dysregulated ribosomal biogenesis may promote an aberrant translational program, facilitating the cancer-driven translation of oncoproteins and potentially contributing to cancer metastasis and therapeutic resistance (76). Therefore, it is important to conduct further studies on the effects of CKM dysregulation resulting from amplifications or mutations in CKM subunits across diverse pathological contexts, with a particular focus on ribosomal biogenesis, cellular growth, and cell proliferation.

## Materials and Methods

Detailed description of the Materials and methods used in this study, including *Drosophila* stocks and maintenance; generation of the transgenic RNAi flies; tagging the endogenous loci of CKM with EGFP or RFP using the CRISPR-Cas9 technique; immunocytochemistry; flow cytometry analyses; single cell RNA-sequencing (scRNA-seq) and subsequent analysis; the HCR RNA-FISH assay; and statistical analyses, are provided in SI Appendix.

## Acknowledgement

This study greatly benefited from the essential contributions of Drs. Jin Sun and Jian-Quan Ni in developing the transgenic RNAi and EGFP-tagged Med12 and Med13 lines. We also thank Dr. Laura Buttitta for sharing the Live Tissue FACS solution recipe and valuable advice, Dr. Robbie Moore at Analytical Cytometry Core of Texas A&M Health Science Center for assistance with flow cytometry analyses, Dr. Andrew Hillhouse at Texas A&M Health Science Center TIGSS Molecular Genomics Core for providing the scRNA-seq service, and the Bloomington *Drosophila* Stock Center (NIH Grant P40OD018537) for supplying fly stocks. The research was supported by a grant from the NIH R01 GM133011 (to J.-Y.J.).

## References

1. S. Hahn, Structure and mechanism of the RNA polymerase II transcription machinery. Nature structural & molecular biology 11, 394–403 (2004).

2. A. C. Schier, D. J. Taatjes, Structure and mechanism of the RNA polymerase II transcription machinery. Genes & development 34, 465–488 (2020).

3. R. G. Roeder, 50+ years of eukaryotic transcription: an expanding universe of factors and mechanisms. Nature structural & molecular biology 26, 783–791 (2019).

4. J. D. Fondell, The Mediator complex in thyroid hormone receptor action. Biochim Biophys Acta 1830, 3867–3875 (2013).

5. S. Malik, R. G. Roeder, The metazoan Mediator co-activator complex as an integrative hub for transcriptional regulation. Nature reviews. Genetics 11, 761–772 (2010).

6. H. M. Bourbon, Comparative genomics supports a deep evolutionary origin for the large, four-module transcriptional mediator complex. Nucleic Acids Res 36, 3993–4008 (2008).

7. H. M. Bourbon et al., A unified nomenclature for protein subunits of mediator complexes linking transcriptional regulators to RNA polymerase II. Mol Cell 14, 553–557 (2004).

8. H. Elmlund et al., The cyclin-dependent kinase 8 module sterically blocks Mediator interactions with RNA polymerase II. Proceedings of the National Academy of Sciences of the United States of America 103, 15788–15793 (2006).

9. K.-L. Tsai et al., A conserved Mediator–CDK8 kinase module association regulates Mediator–RNA polymerase II interaction. 20, 611 (2013).

10. S. Osman et al., The Cdk8 kinase module regulates interaction of the mediator complex with RNA polymerase II. The Journal of biological chemistry 296, 100734 (2021).

11. E. J. Morris et al., E2F1 represses beta-catenin transcription and is antagonized by both pRB and CDK8. Nature 455, 552–556 (2008).

12. J. Zhao, R. Ramos, M. Demma, CDK8 regulates E2F1 transcriptional activity through S375 phosphorylation. Oncogene 32, 3520–3530 (2013).

13. C. J. Fryer, J. B. White, K. A. Jones, Mastermind recruits CycC:CDK8 to phosphorylate the Notch ICD and coordinate activation with turnover. Mol Cell 16, 509–520 (2004).

14. A. J. Donner, S. Szostek, J. M. Hoover, J. M. Espinosa, CDK8 is a stimulus-specific positive coregulator of p53 target genes. Mol Cell 27, 121–133 (2007).

15. C. Alarcon et al., Nuclear CDKs drive Smad transcriptional activation and turnover in BMP and TGF-beta pathways. Cell 139, 757–769 (2009).

16. A. Aleman et al., Mad linker phosphorylations control the intensity and range of the BMP-activity gradient in developing Drosophila tissues. Scientific reports 4, 6927 (2014).

17. X. Zhao et al., Regulation of lipogenesis by cyclin-dependent kinase 8-mediated control of SREBP-1. The Journal of clinical investigation 122, 2417–2427 (2012).

18. J. Bancerek et al., CDK8 Kinase Phosphorylates Transcription Factor STAT1 to Selectively Regulate the Interferon Response. Immunity 38, 250–262 (2013).

19. J. Nemet, B. Jelicic, I. Rubelj, M. J. B. Sopta, The two faces of Cdk8, a positive/negative regulator of transcription. 97, 22–27 (2014).

20. C. B. Fant, D. J. J. T. Taatjes, Regulatory functions of the Mediator kinases CDK8 and CDK19. 10, 76–90 (2019).

21. M. V. Dannappel, D. Sooraj, J. J. Loh, R. J. F. i. c. Firestein, d. biology, Molecular and in vivo functions of the CDK8 and CDK19 kinase modules. 6, 171 (2019).

22. M. T. Knuesel, K. D. Meyer, A. J. Donner, J. M. Espinosa, D. J. Taatjes, The human CDK8 subcomplex is a histone kinase that requires Med12 for activity and can function independently of mediator. Molecular and cellular biology 29, 650–661 (2009).

23. F. Klatt et al., A precisely positioned MED12 activation helix stimulates CDK8 kinase activity. Proceedings of the National Academy of Sciences of the United States of America 117, 2894–2905 (2020).

24. Y. C. Li et al., Structure and noncanonical Cdk8 activation mechanism within an Argonaute-containing Mediator kinase module. Sci Adv 7 (2021).

25. N. Makinen et al., MED12, the mediator complex subunit 12 gene, is mutated at high frequency in uterine leiomyomas. Science 334, 252–255 (2011).

26. M. Turunen et al., Uterine leiomyoma-linked MED12 mutations disrupt mediator-associated CDK activity. Cell reports 7, 654–660 (2014).

27. N. Loncle et al., Distinct roles for Mediator Cdk8 module subunits in Drosophila development. EMBO J 26, 1045–1054 (2007).

28. F. Janody, Z. Martirosyan, A. Benlali, J. E. Treisman, Two subunits of the Drosophila mediator complex act together to control cell affinity. Development 130, 3691–3701 (2003).

29. X. J. Xie et al., CDK8-Cyclin C Mediates Nutritional Regulation of Developmental Transitions through the Ecdysone Receptor in Drosophila. PLoS biology 13, e1002207 (2015).

30. J. Treisman, Drosophila homologues of the transcriptional coactivation complex subunits TRAP240 and TRAP230 are required for identical processes in eye-antennal disc development. Development 128, 603–615 (2001).

31. C. Schiano et al., Involvement of Mediator complex in malignancy. Biochim Biophys Acta 1845, 66–83 (2014).

32. C. Schiano, A. Casamassimi, M. T. Vietri, M. Rienzo, C. Napoli, The roles of mediator complex in cardiovascular diseases. Biochim Biophys Acta 1839, 444–451 (2014).

33. W. Xu, J. Y. Ji, Dysregulation of CDK8 and Cyclin C in tumorigenesis. J Genet Genomics 38, 439–452 (2011).

34. A. D. Clark, M. Oldenbroek, T. G. Boyer, Mediator kinase module and human tumorigenesis. Crit Rev Biochem Mol Biol 50, 393–426 (2015).

35. H.-H. Qiao et al., An efficient and multiple target transgenic RNAi technique with low toxicity in Drosophila. 9, 1–13 (2018).

36. A. C. Groth, M. Fish, R. Nusse, M. P. Calos, Construction of transgenic Drosophila by using the site-specific integrase from phage phiC31. Genetics 166, 1775–1782 (2004).

37. M. Milan, S. Campuzano, A. Garcia-Bellido, Developmental parameters of cell death in the wing disc of Drosophila. Proceedings of the National Academy of Sciences of the United States of America 94, 5691–5696 (1997).

38. X. Li et al., The Mediator CDK8-Cyclin C complex modulates Dpp signaling in Drosophila by stimulating Mad-dependent transcription. PLoS genetics 16, e1008832 (2020).

39. Z. Song, K. McCall, H. Steller, DCP-1, a Drosophila cell death protease essential for development. Science 275, 536–540 (1997).

40. N. Zielke, B. A. Edgar, FUCCI sensors: powerful new tools for analysis of cell proliferation. Wiley Interdiscip Rev Dev Biol 4, 469–487 (2015).

41. L. A. Johnston, B. A. Edgar, Wingless and Notch regulate cell-cycle arrest in the developing Drosophila wing. Nature 394, 82–84 (1998).

42. J. Y. Ji, N. J. Dyson, “Interplay between Cyclin-dependent Kinases and E2F-dependent Transcription” in Cell Cycle Deregulation in Cancer, G. Enders, Ed. (Springer Science 2010), chap. 2, pp. 23–41.

43. D. Kobak, P. Berens, The art of using t-SNE for single-cell transcriptomics. Nat Commun 10, 5416 (2019).

44. B. T. Sherman et al., DAVID: a web server for functional enrichment analysis and functional annotation of gene lists (2021 update). Nucleic Acids Res 50, W216–221 (2022).

45. B. K. Tripathi, K. D. Irvine, The wing imaginal disc. Genetics 220 (2022).

46. M. Ng, F. J. Diaz-Benjumea, J. P. Vincent, J. Wu, S. M. Cohen, Specification of the wing by localized expression of wingless protein. Nature 381, 316–318 (1996).

47. O. Stegle, S. A. Teichmann, J. C. Marioni, Computational and analytical challenges in single-cell transcriptomics. Nat Rev Genet 16, 133–145 (2015).

48. H. M. T. Choi et al., Third-generation in situ hybridization chain reaction: multiplexed, quantitative, sensitive, versatile, robust. Development 145 (2018).

49. R. J. Duronio, P. H. O’Farrell, Developmental control of the G1 to S transition in Drosophila: cyclin Eis a limiting downstream target of E2F. Genes Dev 9, 1456–1468 (1995).

50. T. P. Neufeld, A. F. de la Cruz, L. A. Johnston, B. A. Edgar, Coordination of growth and cell division in the Drosophila wing. Cell 93, 1183–1193 (1998).

51. B. Ren et al., E2F integrates cell cycle progression with DNA repair, replication, and G(2)/M checkpoints. Genes Dev 16, 245–256 (2002).

52. A. M. Anger et al., Structures of the human and Drosophila 80S ribosome. Nature 497, 80–85 (2013).

53. K. Dorner, C. Ruggeri, I. Zemp, U. Kutay, Ribosome biogenesis factors-from names to functions. EMBO J 42, e112699 (2023).

54. U. Rodriguez-Corona, M. Sobol, L. C. Rodriguez-Zapata, P. Hozak, E. Castano, Fibrillarin from Archaea to human. Biol Cell 107, 159–174 (2015).

55. P. L. Monaco, V. Marcel, J. J. Diaz, F. Catez, 2’-O-Methylation of Ribosomal RNA: Towards an Epitranscriptomic Control of Translation? Biomolecules 8 (2018).

56. K. E. Sloan et al., Tuning the ribosome: The influence of rRNA modification on eukaryotic ribosome biogenesis and function. RNA Biol 14, 1138–1152 (2017).

57. D. Tollervey, H. Lehtonen, R. Jansen, H. Kern, E. C. Hurt, Temperature-sensitive mutations demonstrate roles for yeast fibrillarin in pre-rRNA processing, pre-rRNA methylation, and ribosome assembly. Cell 72, 443–457 (1993).

58. T. Gautier, T. Berges, D. Tollervey, E. Hurt, Nucleolar KKE/D repeat proteins Nop56p and Nop58p interact with Nop1p and are required for ribosome biogenesis. Mol Cell Biol 17, 7088–7098 (1997).

59. K. L. Tsai et al., A conserved Mediator-CDK8 kinase module association regulates Mediator-RNA polymerase II interaction. Nature structural & molecular biology 20, 611–619 (2013).

60. E. V. Schneider et al., The Structure of CDK8/CycC Implicates Specificity in the CDK/Cyclin Family and Reveals Interaction with a Deep Pocket Binder. J Mol Biol 412, 251–266 (2011).

61. G. Asher, N. Reuven, Y. Shaul, 20S proteasomes and protein degradation “by default”. Bioessays 28, 844–849 (2006).

62. F. Haque, T. Honjo, N. A. Begum, XLID syndrome gene Med12 promotes Ig isotype switching through chromatin modification and enhancer RNA regulation. Sci Adv 8, eadd1466 (2022).

63. D. C. Stieg et al., A complex molecular switch directs stress-induced cyclin C nuclear release through SCF(Grr1)-mediated degradation of Med13. Mol Biol Cell 29, 363–375 (2018).

64. M. A. Davis et al., The SCF-Fbw7 ubiquitin ligase degrades MED13 and MED13L and regulates CDK8 module association with Mediator. Genes & development 27, 151–156 (2013).

65. D. Y. Youn, A. M. Xiaoli, H. Kwon, F. Yang, J. E. Pessin, The subunit assembly state of the Mediator complex is nutrient-regulated and is dysregulated in a genetic model of insulin resistance and obesity. The Journal of biological chemistry 294, 9076–9083 (2019).

66. J. I. Ovrebo, Y. Ma, B. A. Edgar, Cell growth and the cell cycle: New insights about persistent questions. Bioessays 44, e2200150 (2022).

67. X. Li et al., Cdk8 attenuates lipogenesis by inhibiting SREBP-dependent transcription in Drosophila. Dis Model Mech 15 (2022).

68. P. Tessarz et al., Glutamine methylation in histone H2A is an RNA-polymerase-I-dedicated modification. Nature 505, 564–568 (2014).

69. L. Loza-Muller et al., Fibrillarin methylates H2A in RNA polymerase I trans-active promoters in Brassica oleracea. Front Plant Sci 6, 976 (2015).

70. V. Marcel et al., p53 acts as a safeguard of translational control by regulating fibrillarin and rRNA methylation in cancer. Cancer Cell 24, 318–330 (2013).

71. K. D. Meyer, S. C. Lin, C. Bernecky, Y. Gao, D. J. Taatjes, p53 activates transcription by directing structural shifts in Mediator. Nature structural & molecular biology 17, 753–760 (2010).

72. C. M. Koh et al., Alterations in nucleolar structure and gene expression programs in prostatic neoplasia are driven by the MYC oncogene. Am J Pathol 178, 1824–1834 (2011).

73. J. van Riggelen, A. Yetil, D. W. Felsher, MYC as a regulator of ribosome biogenesis and protein synthesis. Nat Rev Cancer 10, 301–309 (2010).

74. S. S. Grewal, L. Li, A. Orian, R. N. Eisenman, B. A. Edgar, Myc-dependent regulation of ribosomal RNA synthesis during Drosophila development. Nat Cell Biol 7, 295–302 (2005).

75. X. Li, M. Liu, J. Y. Ji, Understanding Obesity as a Risk Factor for Uterine Tumors Using Drosophila. Advances in experimental medicine and biology 1167, 129–155 (2019).

76. A. R. Elhamamsy, B. J. Metge, H. A. Alsheikh, L. A. Shevde, R. S. Samant, Ribosome Biogenesis: A Central Player in Cancer Metastasis and Therapeutic Resistance. Cancer Res 82, 2344–2353 (2022).

