## Supplementary material for "Distinct effects of CDK8 module subunits on cellular growth and proliferation in *Drosophila*": Suppl Methods, Figures, and Tables

**SI Appendix Materials and Methods:**

**Fly strains.** The flies were reared on a standard medium comprising cornmeal, molasses, and yeast, with all genetic crosses maintained at 25˚C. The following strains were obtained from the Bloomington *Drosophila* Stock Center: *ap-Gal4* (BL-3041), *nub-Gal4* (BL-25754), *PCNA-GFP* (*$y^1$ w*; P{w[+mC]=PCNA.ΔNhe::GFP}T110*; BL-25749), and *$w^{1118}$; $Kr^{If-1}$/CyO, P{en1}$wg^{en11}$; P(1)5 P(1)12/TM6B, $Tb^1$* (BL-55124). The recombined line '*ap-Gal4/In(2LR), $wg^{Gla}$, Bc; P(1)5 P(1)12*' was generated in this study.

**Generation of transgenic RNAi flies.** Transgenic RNAi lines enabling the depletion of different combinations of the four CKM subunits were generated using the pNP vector (2). For each gene, pairs of oligoes were synthesized (refer to Table S1 for detailed information on these oligoes). To deplete one gene at a time, the oligoes were annealed and cloned into the pNP vector after digestion with EcoRI/NheI, resulting in plasmids containing one short hairpin. For simultaneous depletion of two genes, a plasmid from the first step was digested with SpeI and ligated with a 300bp short hairpin fragment released from another plasmid via SpeI/XbaI digestion. The resulting plasmids contained two short hairpins. This process was reiterated to generate plasmids containing three or four short hairpins (Table S2). Transgenic fly lines were produced by injecting these constructs into '*y sc v nanos-integrase; attP2'* stock, following the standard procedure (3).

**Tagging the endogenous loci of CKM with EGFP or RFP using the CRISPR-Cas9 technique.** The generation of *CDK8-EGFP* line was previously described (4). To tag RFP to the endogenous locus of *CycC*, three constructs were designed. The pCFD3 vector was used to express two sgRNA to introduce double strand breaks near the 5' of 5' Untranslated Region (UTR) and 3' of 3'UTR of the *CycC* gene. Specifically, primers *gRNA-CycC-L-5.1*, *gRNA-*

*CycC-L-3.1*, *gRNA-CycC-R-5.1*, and *gRNA-CycC-R-3.1* were used, following the general protocol for pCFD3 (referencing http://www.crisprflydesign.org/wp-content/uploads/2014/05/Cloning-with-pCFD3.pdf). All primers and sgRNA sequences are listed in Table S3. The donor DNA contained the *CycC* gene (from the 5' UTR to the stop codon of the gene, primers *CycC-transcript-5.1* and *CycC-transcript-3.1*), *RFP* (primers *CycC-RFP-5.1* and *CycC-RFP-3.1*), the 3' UTR of CycC (*CycC-3UTR-5.1* and *CycC-3UTR-3.1*), 1kb upstream of 5' UTR (*Homology_arm-CycC-L-5.1* and *Homology_arm-CycC-L-3.1*), and 1kb downstream of 3' UTR (*Homology_arm-CycC-R-5.1* and *Homology_arm-CycC-R-3.1*), carried by the pGEM-T Easy vector (Promega, A1360). The Protospacer Adjacent Motif (PAM) sites were mutated to avoid unnecessary cuts, integrated within the aforementioned primers. All three fragments were amplified using PrimeStar Max premix (Takara, R045A) and assembled into the pGEM-T Easy vector using NEBuilder HiFi DNA Assembly (NEB, E5520S). Subsequently, all three constructs were co-injected into the embryos of the fly strain *nos-Cas9* by the Rainbow Transgenic Flies, Inc (Camarillo, CA).

To generate the *Med12-EGFP* and *Med13-EGFP* strains, the homologous arms of *Med12* or *Med13* from *Drosophila* genomic DNA were amplified and subsequently cloned into the *Avr*II and *Spe*I sites of the HP1a-4XP3-mCherry donor vector (5) using Gibson assembly. Next, an EGFP tag was cloned into the C-terminus of Med12 or Med13, positioned immediately upstream of the translational stop codon. This process resulted in the construction of the *Med12-EGFP* and *Med13-EGFP* donor plasmids. The design and cloning of sgRNAs into the U6b-sgRNA-short vector were performed as previously described (6). Finally, the donor vector along with the corresponding sgRNA vector were injected into *P{nos-Cas9}attP40* fly embryos using established protocols (5).

**Immunocytochemistry.** The immunostaining of wing discs followed a protocol similar to that described previously (7). Specifically, phosphorylated Histone 3 Serine 10 (PH3) was detected using an anti-PH3 monoclonal antibody (diluted 1:500 in PBS-Triton X-100-NGS-BSA; obtained from the Sigma-Aldrich). Cleaved *Drosophila* Dcp-1 was detected using anti-cleaved Dcp-1 (Asp215) antibody (1:500 diluted in PBS-Triton X-100-NGS-BSA; Cell Signaling, #9578). Fibrillarin/Fib was detected by anti-Fib (diluted 1:500 in PBS-Triton X-100-NGS-BSA; Invitrogen, #MA3-16771). Secondary antibodies with fluorophores were used: goat anti-guinea pig (106-545-003), goat anti-mouse (115-545-003), or goat anti-rabbit (111-545-003), all obtained from Jackson Immunological Laboratories.

**Flow cytometry analyses.** twenty wing imaginal discs were dissected from third instar larvae in PBS. PBS was then replaced with 500 μL of Live Tissue FACS solution (comprising 900 μL 0.25% Trypsin-EDTA from ThermoFisher, 100 μL 10X PBS to achieve a final concentration of 1X, and 0.5 μL of Dye Cycle Violet) for simultaneous cell dissociation and DNA staining, incubated at 37 ˚C for 15min (8). Subsequently, the cell suspension was cooled on ice and directly sorted using a Fortessa X-20. Excitation of Dye Cycle Violet was achieved with a 405 nm laser, while detection of GFP signal used a 488 nm laser. The collected data were processed and analyzed using FCS Express 7. GFP-positive cells and those exhibiting 2N/4N DNA content were selected for cell size estimation and cell cycle analysis. The "Cycle 1, Model 1" algorithm from FCS Express 7 was applied for cell cycle model fitting.

**Single cell RNA-sequencing (scRNA-seq) and subsequent analysis.** 100-120 wing imaginal discs were dissected and dissociated in 500 μL 0.25% Trypsin-EDTA (ThermoFisher) at 37°C for 10 min. The resulting cell suspension underwent two washes with PBS and was then passed through a 35 μm filter (9). The filtered cells were resuspended and sent to Texas A&M Health

Science Center TIGSS Molecular Genomics Core for 10X Genomics library prep and subsequential sequencing. The r6.35 release of the *Drosophila* genome, obtained from Flybase, served as the reference, with a manually added GFP sequence. Alignment was performed using CellRanger 3.0.2, followed by further processing of the count data with Seurat. The scripts used in this analysis will be available at a Github link upon acceptance. Gene lists generated by Seurat were subsequently enriched and plotted using clusterProfiler (10).

**The HCR RNA-FISH assay.** The multiplexed HCR analyses were conducted as described previously (11). Molecular Instruments supplied the probe sets and amplifiers: B1-Alexa Fluor 488 amplifiers were paired with the probe set designed for *CycE* (lot number: RTE467; GenBank Accession #: NM_057611.5), B2-Alexa Fluor 594 amplifiers were used with the probe sets designed for *Mcm5* (lot number: RTE468; GenBank: NM_079584.4), and B3-Alexa Fluor 647 amplifiers were used with the probe sets designed for *stg* (lot number: RTD617; GenBank: NM_079823.4). Confocal imaging was performed using a Zeiss LSM900 confocal microscope system, and images were processed using Adobe Photoshop 2021.

**Statistical analyses.** In this study, p-values were calculated using one-tailed unpaired *t*-tests, while error bars showed in the figures represent the standard deviation. Experiments in this study are based on a minimum of three independent biological replicates. Significance levels were indicated as follows: */# for $p<0.05$, **/## for $p<0.01$, and ***/### for $p<0.001$.

**References:**

1. M. Vaskova *et al.*, Genetic analysis of the Drosophila 63F early puff. Characterization of mutations in E63-1 and maggie, a putative Tom22. *Genetics* **156**, 229-244 (2000).

2. H. H. Qiao *et al.*, An efficient and multiple target transgenic RNAi technique with low toxicity in Drosophila. *Nat Commun* **9**, 4160 (2018).

3. J. Q. Ni *et al.*, A genome-scale shRNA resource for transgenic RNAi in Drosophila. *Nat Methods* **8**, 405-407 (2011).

4. M. Liu, Xie, X.-J., Li, X., Ren, X., Sun, J., Lin, Z., Hemba-Waduge, R., and Ji, J.Y., Transcriptional coupling of telomeric retrotransposons with the cell cycle. *https://doi.org/10.1101/2023.09.30.560321* (2024).

5. X. Ren *et al.*, Enhanced specificity and efficiency of the CRISPR/Cas9 system with optimized sgRNA parameters in Drosophila. *Cell Rep* **9**, 1151-1162 (2014).

6. X. Ren *et al.*, Optimized gene editing technology for Drosophila melanogaster using germ line-specific Cas9. *Proceedings of the National Academy of Sciences of the United States of America* **110**, 19012-19017 (2013).

7. X. Li *et al.*, The Mediator CDK8-Cyclin C complex modulates Dpp signaling in Drosophila by stimulating Mad-dependent transcription. *PLoS genetics* **16**, e1008832 (2020).

8. A. F. de la Cruz, B. A. Edgar, Flow cytometric analysis of Drosophila cells. *Methods Mol Biol* **420**, 373-389 (2008).

9. M. Deng *et al.*, Single cell transcriptomic landscapes of pattern formation, proliferation and growth in Drosophila wing imaginal discs. *Development* **146** (2019).

10. G. Yu, L. G. Wang, Y. Han, Q. Y. He, clusterProfiler: an R package for comparing biological themes among gene clusters. *Omics : a journal of integrative biology* **16**, 284-287 (2012).

11. X. Li *et al.*, Cdk8 attenuates lipogenesis by inhibiting SREBP-dependent transcription in Drosophila. *Dis Model Mech* **15** (2022).

**The Supplementary Figures and Tables**

**Fig. S1 The effect of depleting CKM subunits using *ap-Gal4* on the wing blade and thorax.** Representative adult females of the following genotypes are shown: (A) *ap-Gal4>UAS-2XGFP/+*; (B) *ap-Gal4>UAS-2XGFP/+; UAS-Cdk8-RNAi CycC-RNAi/+*; (C) *ap-Gal4>UAS-2XGFP/+; UAS-Med12-RNAi/+* and (D) *ap-Gal>UAS-2XGFP/+; UAS-Cdk8-RNAi CycC-RNAi Med12-RNAi/+*. Additionally, representative dead females dissected out from the pupal case are shown for: (E) *ap-Gal4>UAS-2XGFP/+; UAS-Med12-RNAi Med13-RNAi/+*, and (F) *ap-Gal4>UAS-2XGFP/+; UAS-Cdk8-RNAi CycC-RNAi Med12-RNAi Med13-RNAi/+*.

**Fig. S2 Depleting CKM subunits had little effects on apoptosis in wing discs.** Representative confocal images focused on the wing pouch area of wing discs stained with DAPI (blue) and anti-Dcp-1 (red) antibody are provided for the following genotypes: (A) *ap-Gal4>UAS-2XGFP/+*; (B) *ap-Gal4>UAS-2XGFP/+; UAS-Med12-RNAi Med13-RNAi/+*; and (C) *ap-Gal4>UAS-2XGFP/+; UAS-Cdk8-RNAi CycC-RNAi Med12-RNAi Med13-RNAi/+*. The scale bar in panel C: 25μm.

**Fig. S3 The effect of depleting the CKM subunits on cell-cycle progression, assessed using the Fly-FUCCI system.** Representative confocal images focused on the wing pouch area of wing discs are provided for the following genotypes: (A) *nub-Gal4/+; UAS-GFP-E2f1*$_{1-230}$*, nls-mRFP-CycB*$_{1-266}$*/+* with GFP-E2f1$_{1-230}$ (green) and nls-mRFP-CycB$_{1-266}$ (red); (B) *nub-Gal4/+; UAS-GFP-E2f1*$_{1-230}$*, nls-mRFP-CycB*$_{1-266}$*/UAS-Cdk8-RNAi CycC-RNAi*; (C) *nub-Gal4/+; UAS-GFP-E2f1*$_{1-230}$*, nls-mRFP-CycB*$_{1-266}$*/UAS-Med12-RNAi*; (D) *nub-Gal4/+; UAS-GFP-E2f1*$_{1-230}$*, nls-mRFP-CycB*$_{1-266}$*/UAS-Cdk8-RNAi CycC-RNAi Med12-RNAi*; (E) *nub-Gal4/+; UAS-GFP-E2f1*$_{1-230}$*, nls-mRFP-CycB*$_{1-266}$*/UAS-Med12-RNAi Med13-RNAi*; (F) *nub-Gal4/+; UAS-GFP-E2f1*$_{1-230}$*, nls-mRFP-CycB*$_{1-266}$*/UAS-Cdk8-RNAi CycC-RNAi Med12-RNAi Med13-RNAi*. (G)

Quantification of the relative number of green, red, or yellow pixels. Three wing discs of each genotype were used for quantification. The scale bar in panel F: 25µm. */#/$ $p<0.05$; **/##/$$ $p<0.01$; ***/###/$$$ $p<0.001$.

**Fig. S4 The cell cycle model fit of DNA content in cells from the wing disc with CKM depletion.** Histogram of cell-cycle model fit of DyeCycle Violet signal of the cells are shown for the following genotypes: (A) *nub-Gal4/+; UAS-mCD8::GFP/*;+; (B) *nub-Gal4/+; UAS-mCD8::GFP/UAS-Cdk8-RNAi CycC-RNAi* (red); (C) *nub-Gal4/+; UAS-mCD8::GFP/UAS-Med12-RNAi*; (D) *nub-Gal4/+; UAS-mCD8::GFP/UAS-Cdk8-RNAi CycC-RNAi Med12-RNAi*; (E) *nub-Gal4/+; UAS-mCD8::GFP/UAS-Med12-RNAi Med13-RNAi*; and (F) *nub-Gal4/+; UAS-mCD8::GFP/UAS-Cdk8-RNAi CycC-RNAi Med12-RNAi Med13-RNAi*. (G) Quantification and statistical analysis of cell-cycle phases using model fitting are also provided, including %G1 (G1 phase cell population), %G2 (G2 phase cell population), %S (S phase cell population), Chi sq (Chi-square value of the model fitting), and B.A.D. (Background Aggregates and Debris).

**Fig. S5 The expression distribution of landmark genes in tSNE maps containing all the detected cells.** (A) Expression distribution of *nubbin*; (B) Expression distribution of *teashirt*; and (C) Expression distribution of *pannier*.

**Fig. S6 The scRNA-seq analyses of wing disc cell type populations from wing discs with depletion of different CMK subunits.** (A) tSNE map for *ap-Gal4>UAS-2XGFP/+; +*; (B) tSNE map for *ap-Gal4>UAS-2XGFP/+; UAS-Cdk8-RNAi CycC-RNAi/+*; (C) tSNE map for *ap-Gal4>UAS-2XGFP/+; UAS-Med12-RNAi/+*; and (D) tSNE map for *ap-Gal>UAS-2XGFP/+; UAS-Cdk8-RNAi CycC-RNAi Med12-RNAi/+*. (E) The cell population of each cluster of genotypes is color-coded for reference.

**Fig. S7 Venn diagrams comparing differentially expressed genes in *nub*-positive cells of different genotypes.** Gene expression levels that are higher in dorsal compartment (GFP+) compared to the ventral compartment (GFP-) are shown for the following genotypes: (A) *ap-Gal4>UAS-2XGFP/+* and *ap-Gal4>UAS-2XGFP/+; UAS-Cdk8-RNAi CycC-RNAi/+*; (B) *ap-Gal4>UAS-2XGFP/+* and *ap-Gal4>UAS-2XGFP/+; UAS-Med12-RNAi/+*; and (C) *ap-Gal4>UAS-2XGFP/+* and *ap-Gal>UAS-2XGFP/+; UAS-Cdk8-RNAi CycC-RNAi Med12-RNAi/+*. Gene expression levels that are lower in the dorsal compartment (GFP+) compared to the ventral compartment (GFP-) are shown for: (D) *ap-Gal4>UAS-2XGFP/+* and *ap-Gal4>UAS-2XGFP/+; UAS-Cdk8-RNAi CycC-RNAi/+*; (E) *ap-Gal4>UAS-2XGFP/+* and *ap-Gal4>UAS-2XGFP/+; UAS-Med12-RNAi/+*; and (F) *ap-Gal4>UAS-2XGFP/+* and *ap-Gal>UAS-2XGFP/+; UAS-Cdk8-RNAi CycC-RNAi Med12-RNAi/+*.

**Fig. S8 The role of CKM in regulating the expression of E2F1 target gene *stg*.** Confocal images illustrating the expression of BFP (A-C) and mRNA transcripts of *stg* (magenta) (A'-C' and D-F) within the wing pouch region of wing discs. Genotypes: (A/A') *en-Gal4/+; UAS-BFP/+* (control); (B/B') *en-Gal4/+; UAS-Cdk8-RNAi CycC-RNAi/UAS-BFP*; (C/C') *en-Gal4/+; UAS-Med12-RNAi/UAS-BFP*; (D) *en-Gal4/+; UAS-*$Dp^{RNAi}$*/UAS-BFP*; (E) *en-Gal4/+; UAS-*$Rbf1^{RNAi}$*/UAS-BFP*; and (F) *en-Gal4/+; UAS-*$CycE^{+}$*/UAS-BFP*. 'A/P' indicates the anterior-posterior compartment boundary. The scale bar in panel C: 20 μm.

**Table S1. Primer sequences utilized to generate pNP vectors for CKM subunits.**

**Table S2. List of the transgenic RNAi lines for CKM subunits.**

**Table S3. Primers and sgRNAs utilized to generate EGFP- or RFP-tagged CKM subunits.**

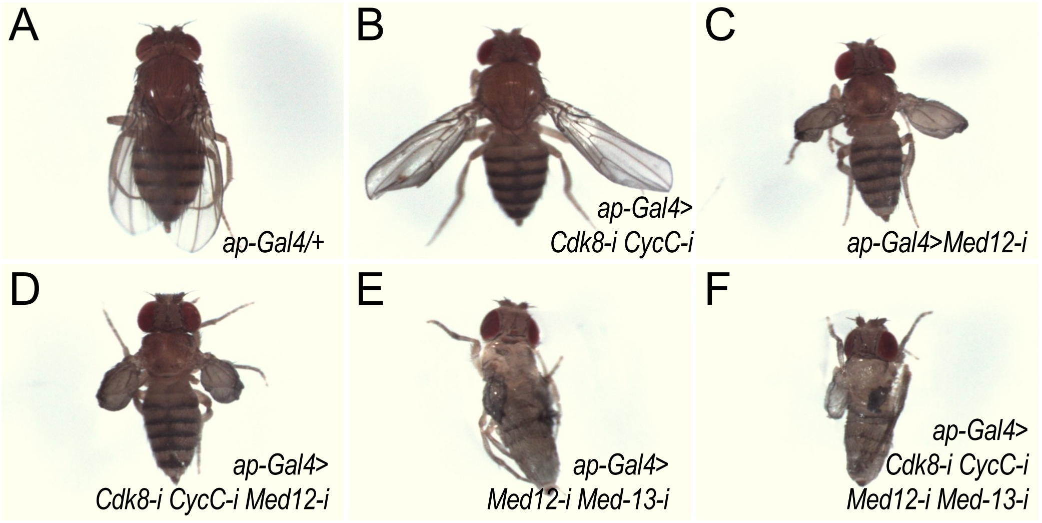


**Fig. S1**

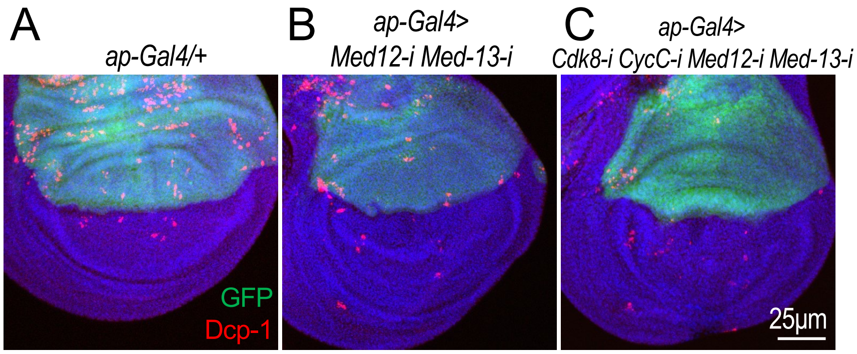


**Fig. S2**

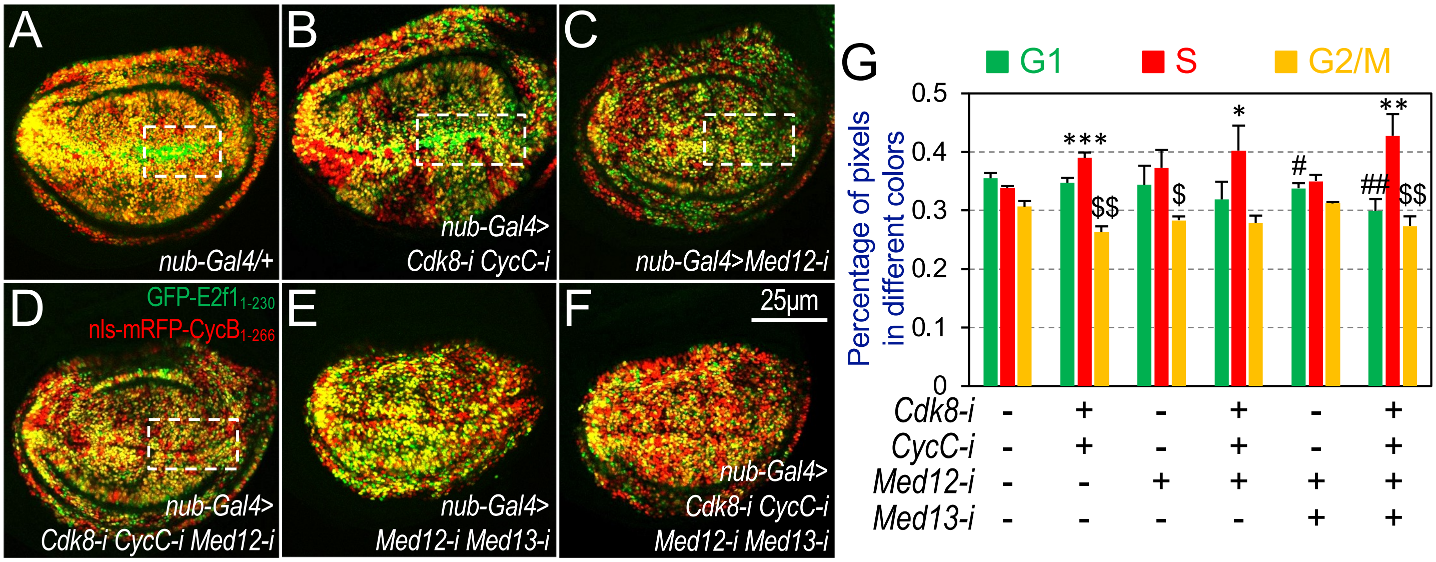


**Fig. S3**

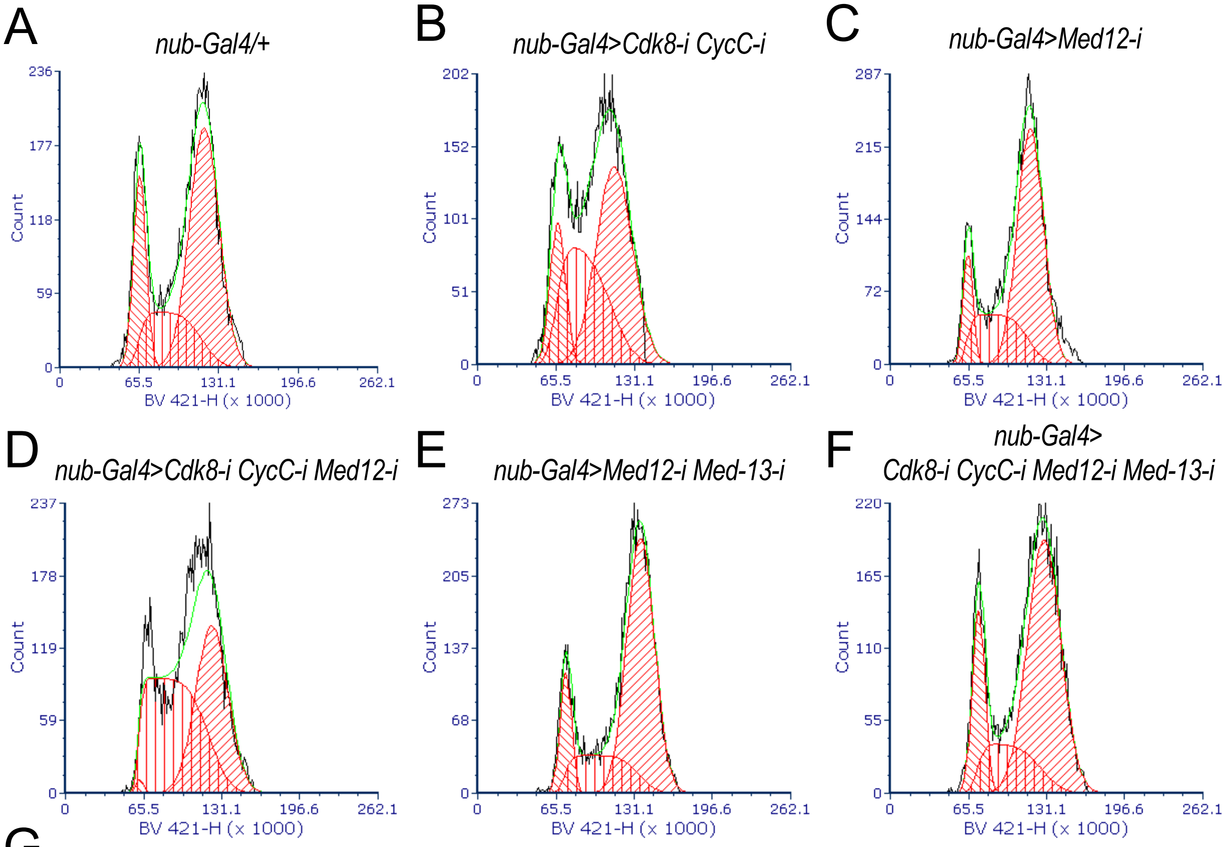


G

| | % G1 | % G2 | % S | Total cell number | Chi sq | B.A.D |
|---|---|---|---|---|---|---|
| *nub-Gal4/+* | 19.88 | 57.59 | 22.73 | 10185 | 1.61 | 0 |
| *nub-Gal4>Cdk8-i CycC-i* | 14.97 | 49.07 | 35.97 | 10357 | 1.43 | 0 |
| *nub-Gal4>Med12-i* | 11.21 | 63.56 | 25.23 | 10366 | 1.28 | 0 |
| *nub-Gal4>Cdk8-i CycC-i Med12-i* | 0.89 | 45.29 | 53.82 | 10602 | 4.99 | 0 |
| *nub-Gal4>Med12-i Med-13-i* | 12.99 | 66.19 | 20.81 | 10507 | 1.43 | 0 |
| *nub-Gal4>Cdk8-i CycC-i Med12-i Med-13-i* | 17.35 | 63.81 | 18.84 | 10521 | 1.44 | 0 |

**Fig. S4**

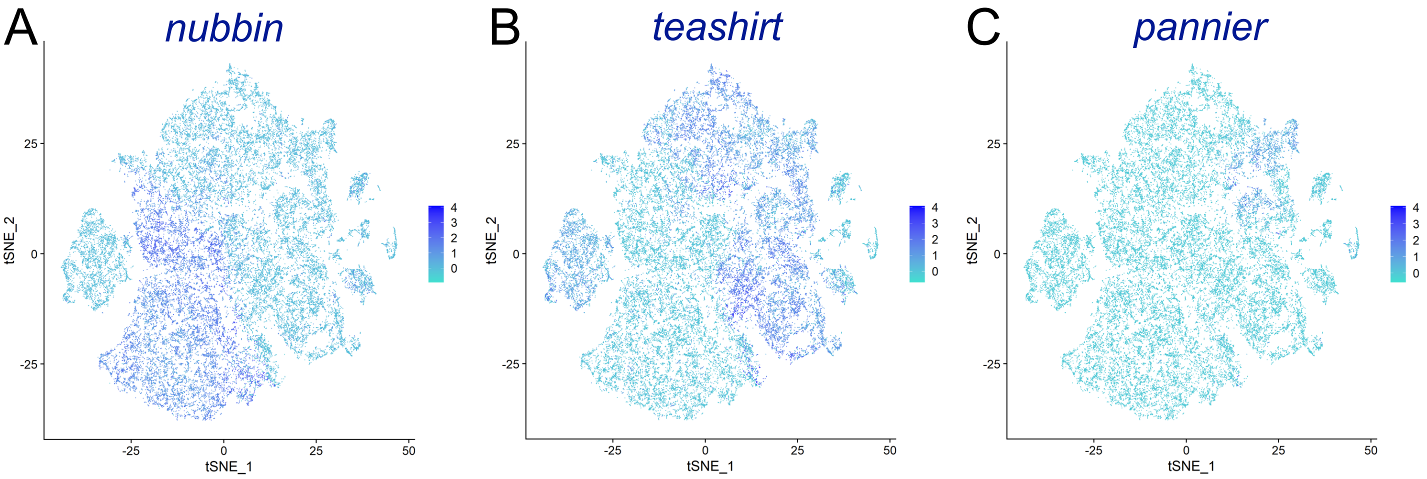


**Fig. S5**

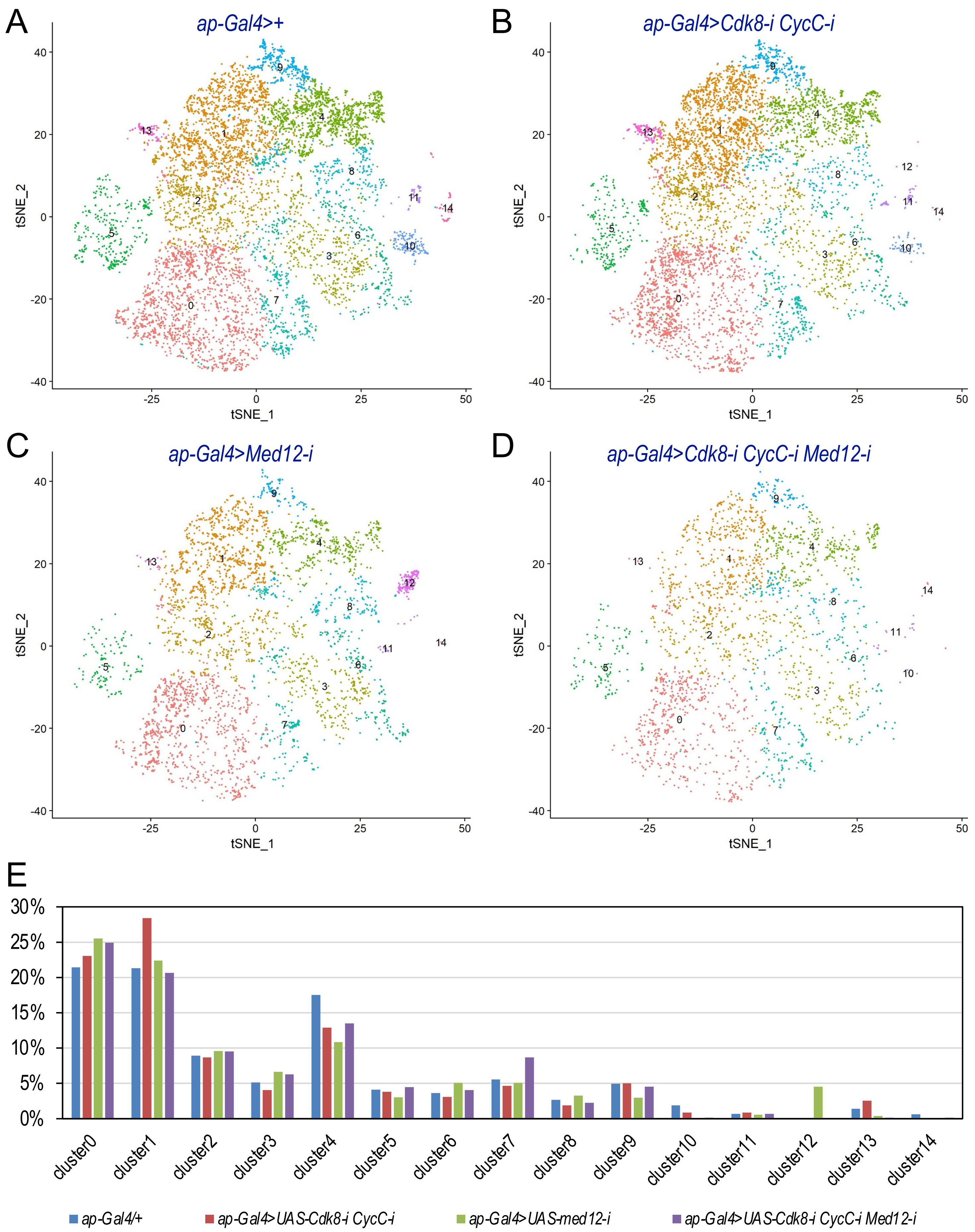
A
ap-Gal4>+
B
ap-Gal4>Cdk8-i CycC-i
C
ap-Gal4>Med12-i
D
ap-Gal4>Cdk8-i CycC-i Med12-i
tSNE_1
tSNE_2
E
cluster0
cluster1
cluster2
cluster3
cluster4
cluster5
cluster6
cluster7
cluster8
cluster9
cluster10
cluster11
cluster12
cluster13
cluster14
ap-Gal4/+
ap-Gal4>UAS-Cdk8-i CycC-i
ap-Gal4>UAS-med12-i
ap-Gal4>UAS-Cdk8-i CycC-i Med12-i

**Fig. S6**

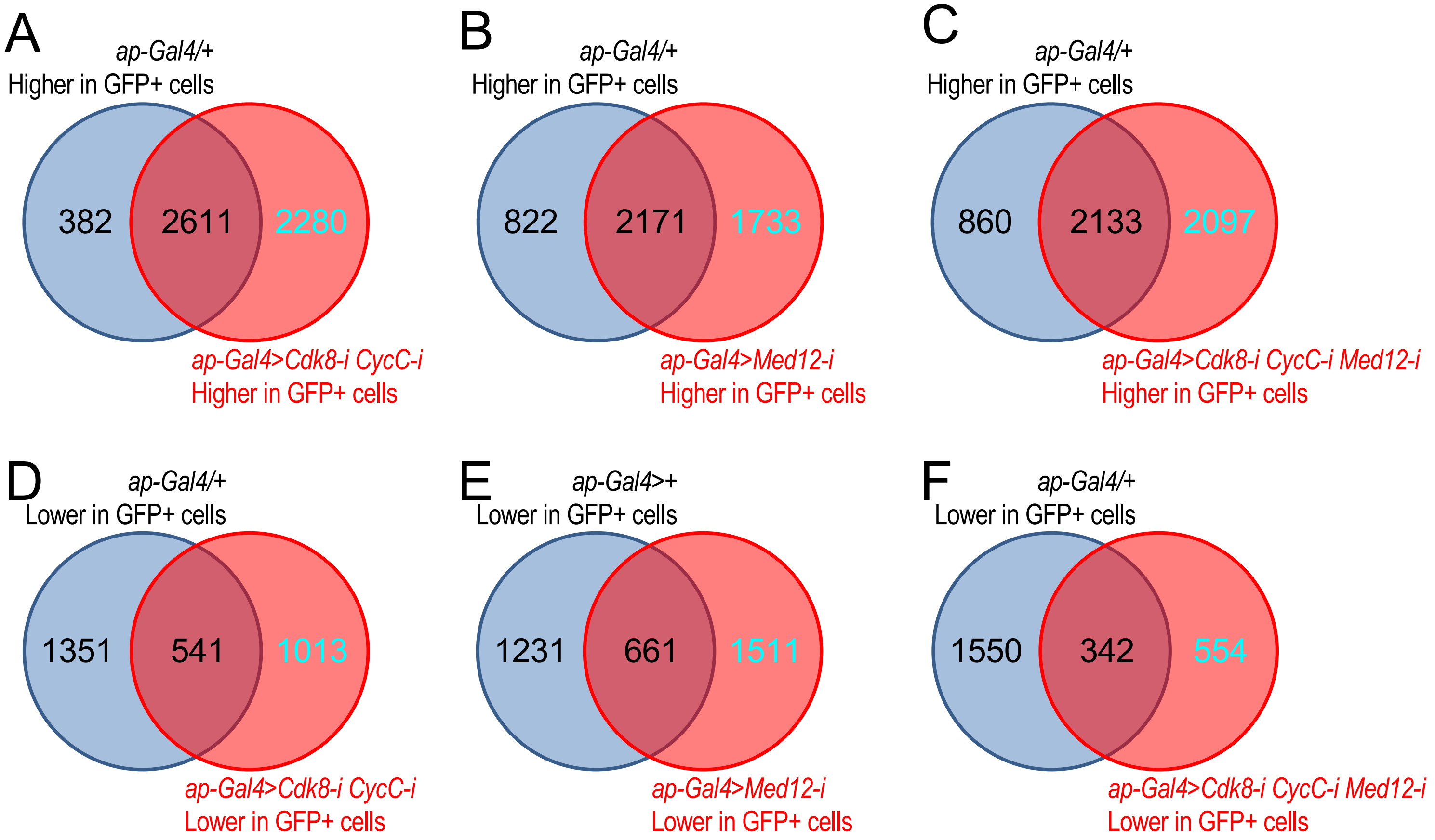
A
ap-Gal4/+
Higher in GFP+ cells
382
2611
2280
ap-Gal4>Cdk8-i CycC-i
Higher in GFP+ cells
B
ap-Gal4/+
Higher in GFP+ cells
822
2171
1733
ap-Gal4>Med12-i
Higher in GFP+ cells
C
ap-Gal4/+
Higher in GFP+ cells
860
2133
2097
ap-Gal4>Cdk8-i CycC-i Med12-i
Higher in GFP+ cells
D
ap-Gal4/+
Lower in GFP+ cells
1351
541
1013
ap-Gal4>Cdk8-i CycC-i
Lower in GFP+ cells
E
ap-Gal4>+
Lower in GFP+ cells
1231
661
1511
ap-Gal4>Med12-i
Lower in GFP+ cells
F
ap-Gal4/+
Lower in GFP+ cells
1550
342
554
ap-Gal4>Cdk8-i CycC-i Med12-i
Lower in GFP+ cells

**Fig. S7**

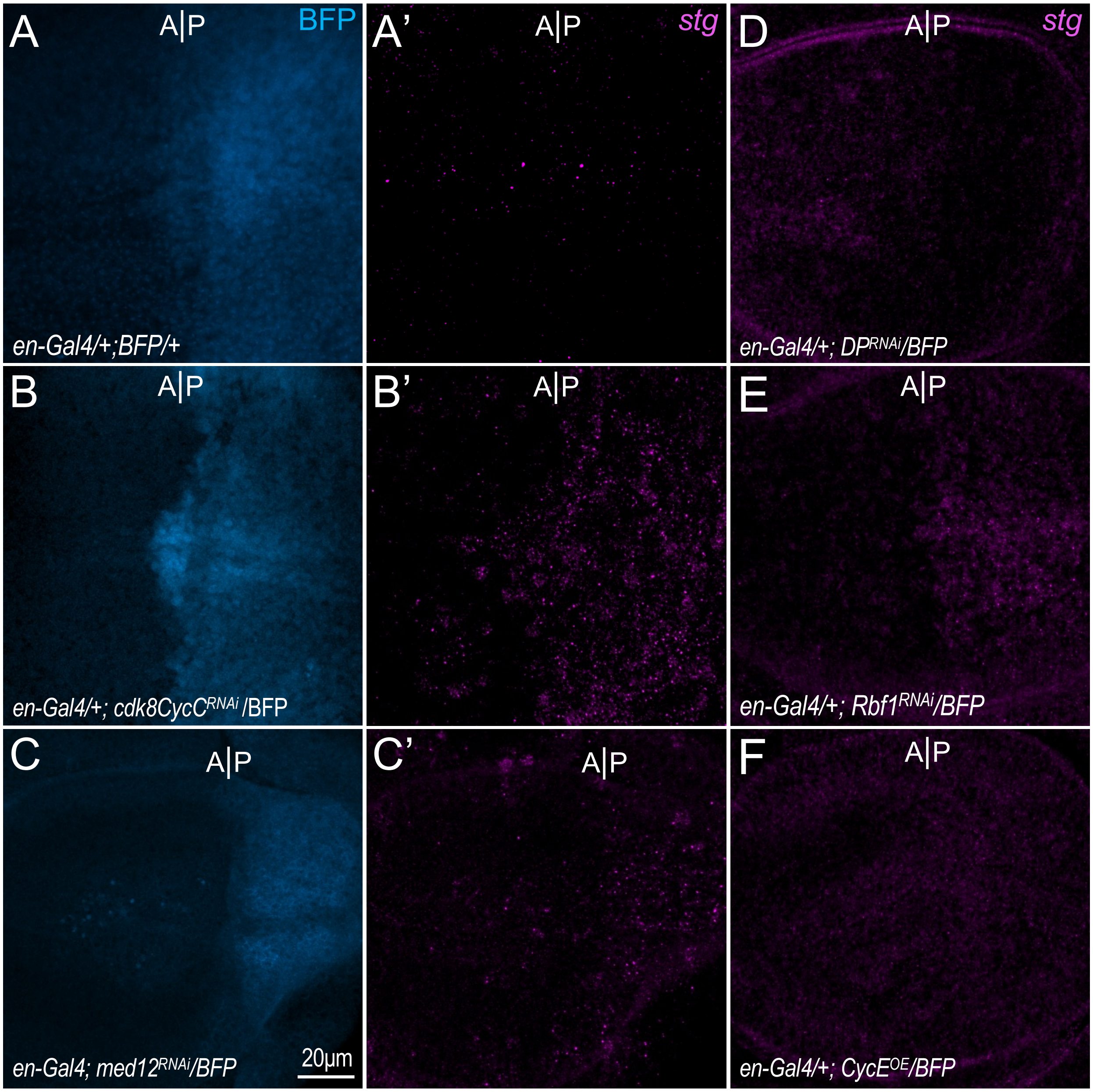
A
A|P
BFP
en-Gal4/+;BFP/+
A'
A|P
stg
D
A|P
stg
en-Gal4/+; DPRNAi/BFP
B
A|P
en-Gal4/+; cdk8CycCRNAi /BFP
B'
A|P
E
A|P
en-Gal4/+; Rbf1RNAi/BFP
C
A|P
en-Gal4; med12RNAi/BFP
20µm
C'
A|P
F
A|P
en-Gal4/+; CycEOE/BFP

**Fig. S8**

**Table S1. Primer sequences used to generate pNP vectors for CKM subunits.**

| Genes | Oligo sequences |
|---|---|
| *Cdk8* (forward) | ctagcagtCAAGGTGTTCCTGTTGATCGAtagttatattcaagcataTCGATCAACAGGAACACCTTGgcg |
| *Cdk8* (reverse) | aattcgcCAAGGTGTTCCTGTTGATCGAtatgcttgaatataactaTCGATCAACAGGAACACCTTGactg |
| *CycC* (forward) | ctagcagtCACGGCGACGGTTTACTTCAAtagttatattcaagcataTTGAAGTAAACCGTCGCCGTGgcg |
| *CycC* (reverse) | aattcgcCACGGCGACGGTTTACTTCAAtatgcttgaatataactaTTGAAGTAAACCGTCGCCGTGactg |
| *Med12* (forward) | ctagcagtCAGATGCTACGACAACGACAAtagttatattcaagcataTTGTCGTTGTCGTAGCATCTGgcg |
| *Med12* (reverse) | aattcgcCAGATGCTACGACAACGACAAtatgcttgaatataactaTTGTCGTTGTCGTAGCATCTGactg |
| *Med13* (forward) | ctagcagtGAtGGAGACtAACGAAAtAAAtagttatattcaagcataTTTATTTCGTTAGTCTCCATCgcg |
| *Med13* (reverse) | aattcgcGAtGGAGACtAACGAAAtAAAtatgcttgaatataactaTTTATTTCGTTAGTCTCCATCactg |

**Table S2 List of the transgenic RNAi lines for CKM subunits and stocks from the Bloomington stock center.**

| Tsinghua Fly Center ID | Gene Symbol | Vector | Hairpin Location | Discription |
|---|---|---|---|---|
| TH00494.N | *Cdk8* | pNP | *attP2* | |
| TH00496.N | *Med13* | pNP | *attP2* | |
| TH00498.N | *CycC* | pNP | *attP2* | |
| TH00500.N | *Med12* | pNP | *attP2* | |
| TH00502.N | *CycC, Med13* | pNP | *attP2* | The hairpin from TH00496.N is cloned into the SpeI site of TH00498.N |
| TH00503.N | *CycC, Med12* | pNP | *attP2* | The hairpin from TH00500.N is cloned into the SpeI site of TH00498.N |
| TH00504.N | *CycC, Cdk8* | pNP | *attP2* | The hairpin from TH00494.N is cloned into the SpeI site of TH00498.N |
| TH00508.N | *Med13, Med12* | pNP | *attP2* | The hairpin from TH00500.N is cloned into the SpeI site of TH00496.N |
| TH00509.N | *Med13, Cdk8* | pNP | *attP2* | The hairpin from TH00494.N is cloned into the SpeI site of TH00496.N |
| TH00510.N | *Med12, Cdk8* | pNP | *attP2* | The hairpin from TH00494.N is cloned into the SpeI site of TH00500.N |
| TH00595.S | *Cdk8, Med12* | pNP | *attP2* | The hairpin from TH00500.N is cloned into the SpeI site of TH00494.N |
| TH00596.S | *Cdk8, Med13* | pNP | *attP2* | The hairpin from TH00496.N is cloned into the SpeI site of TH00494.N |
| TH00597.S | *Med12, CycC* | pNP | *attP2* | The hairpin from TH00498.N is cloned into the SpeI site of TH00500.N |
| TH00598.S | *Med13, CycC* | pNP | *attP2* | The hairpin from TH00498.N is cloned into the SpeI site of TH00496.N |
| TH00599.S | *CycC, Cdk8, Med12* | pNP | *attP2* | The hairpin from TH00500.N is cloned into the SpeI site of TH00504.N |
| TH00600.S | *CycC, Cdk8, Med13* | pNP | *attP2* | The hairpin from TH00496.N is cloned into the SpeI site of TH00504.N |
| TH00601.S | *Med13, Med12, CycC* | pNP | *attP2* | The hairpin from TH00498.N is cloned into the SpeI site of TH00508.N |
| TH00602.S | *Med13, Med12, Cdk8* | pNP | *attP2* | The hairpin from TH00494.N is cloned into the SpeI site of TH00508.N |
| TH00605.S | *Med13, Med12, Cdk8, CycC* | pNP | *attP2* | The hairpin from TH00504.N is cloned into the SpeI site of TH00508.N |

| BDSC Stock # | Genotypes | Comments |
|---|---|---|
| 30515 | *y[1] v[1]; P{y[+t7.7] v[+t1.8]=TRiP.JF02519}attP2* | *UAS-Dp-RNAi* |
| 36744 | *y[1] sc[*] v[1] sev[21]; P{y[+t7.7] v[+t1.8]=TRiP.HMS03004}attP2/TM3, Sb[1]* | *UAS-Rbf1-RNAi* |
| 4781 | *w[*]; P{w[+mC]=UAS-CycE.L}ML1* | *UAS-CycE-OE* |

**Table S3. Primers and sgRNAs utilized to generate EGFP- or RFP-tagged CKM subunits.**

| Primer name | Primer sequence |
|---|---|
| *Homology_arm-CycC-L-5.1* | GGCCGCCATGGCCGCGGGATCCCTGATAGGCGGTGGCGTG |
| *Homology_arm-CycC-L-3.1* | AGCTGATTAGGTCGGCCAATATTTTCGTGTTAGCTATCCA |
| *Homology_arm-CycC-R-5.1* | TTTATAATAATTATTTCTAAATTAAATTACATAATTGCTT |
| *Homology_arm-CycC-R-3.1* | CAGGCGGCCGCACTAGTGATTCCAAAAAGAAGAAGTCGAA |
| *CycC-transcript-5.1* | TGGATAGCTAACACGAAAATATTGGCCGACCTAATCAGCT |
| *CycC-transcript-3.1* | ACATTCTTGGAAGACCTCATACGCTGAGGCGGTGGTTTCG |
| *CycC-RFP-5.1* | CGAAACCACCGCCTCAGCGTATGAGGTCTTCCAAGAATGT |
| *CycC-RFP-3.1* | ATGATTATGAAGCTCTTCTATTAAAGGAACAGATGGTGGC |
| *CycC-3UTR-5.1* | GCCACCATCTGTTCCTTTAATAGAAGAGCTTCATAATCAT |
| *CycC-3UTR-3.1* | AAGCAATTATGTAATTTAATTTAGAAATAATTATTATAAA |
| *gRNA-CycC-L-5.1* | gtcgTTTAGAAATAATTATTATAA |
| *gRNA-CycC-L-3.1* | aaacTTATAATAATTATTTCTAAA |
| *gRNA-CycC-R-5.1* | gtcgATTGGCCGACCTAATCAGCT |
| *gRNA-CycC-R-3.1* | aaacAGCTGATTAGGTCGGCCAAT |
| *Med12-left-F* | GCGAATTGGGTACAAGCTCCTAGGGTCGCAGAATCCGATGATGC |
| *med12-left-R* | GGTACCATACAAGCTTGTACGGTGGTGGTTGATACTG |
| *med12-right-F* | GTACAAGCTTGTATGGTACCGAAGACACTTCTTAATCGTAAGCC |
| *med12-right-R* | GAACAAAAGCTGGAGCTCACTAGTGAACATGGGAAAGAGATCTGGG |
| *Med12-sgRNA* | GTGTCTTCGATCTAGTACGG |
| *med13-left-F* | GCGAATTGGGTACAAGCTCCTAGGTGCTCAGCAAGCAACAGCTG |
| *med13-left-R* | GGTACCATACAAGCTTGGCTATTGCAGCCGTCAGAT |
| *med13-right-F* | AGCCAAGCTTGTATGGTACCGGTCCGGGATCTCAGATCCT |
| *med13-right-R* | GAACAAAAGCTGGAGCTCACTAGTACTCTTCCTAACCGCTCATG |
| *Med13-sgRNA* | GCTGCAATAGCCTAAGGTCC |
| *eGFP-F* | TTAAGCTTGTGAGCAAGGGCGAGGAGC |
| *eGFP-R* | TTGGTACCTACTTGTACAGCTCGTCCAT |
